# Reservoir water in Singapore contains ESBL-producing and carbapenem-resistant bacteria with conjugatable conserved gene cluster transfer between different species

**DOI:** 10.1101/2021.06.13.448270

**Authors:** Yang Zhong, Siyao Guo, Joergen Schlundt

## Abstract

As the role of the aquatic environment in the “One Health” approach has called increasing attention, the studies of Antimicrobial resistance (AMR) spreading in the water bodies have been reported worldwide. However, there are still limited studies on the AMR carrier in the reservoir water in Singapore. Since 2018, our group has collect water samples from six reservoirs in Singapore and isolated the beta-lactam resistant bacteria from them. We then characterized the isolates with Whole-genome sequencing (WGS) and successfully identified ESBL-producing bacteria from three sampling reservoirs, and confirmed their resistance with both phenotypic and sequencing methods. To better understand the AMR spreading locally, we compared our isolates with isolates from other WGS studies in Singapore covered humans, food, and the enviroment. From there, we noticed the same sequence type (ST) as ST10, ST23, and ST38 has been shared among the environment, food, and humans, as well as the same beta-lactamase genes, are widely distributed among multiple sources. Further genetic environment comparison of beta-lactamase has suggested their spreading as conserved gene clusters among different species and sources. And this hypothesis has been supported by the successful conjugation of *bla*_CTX-M-15_ from *Klebsiella pneumonia* to *Escherichia coli (E .coli)*. We also applied the shotgun metagenomic sequencing to understand the community of bacteria in reservoir water and detect the AMR genes. The composition of bacteria has shown different diversity among different samples. Besides, different beta-lactamase genes have been identified compared to culture depended methods. Here, we suggest that sequencing analysis has great potential in understanding AMR spreading in the “One-Health” approach. A genetic-based AMR risk assessment is in urgent need in Singapore.

## Introduction

As stated by the World Health Organisation (WHO), by 2050, antimicrobial resistance (AMR) will result in an extra burden of 1.2 trillion USD on health expenditure every year(Adeyi, Baris et al. 2017). Significantly, the AMR problem has become a serious challenge to public health, and the concert on AMR from food and the environment to humans has called increasing attention(FAO- OIE-WHO 2010). Based on the WHO report “Antimicrobial Resistance: An Emerging Water, Sanitation and Hygiene Issue,” the aquatic environment can perform like a reservoir of AMR bacteria from both humans and animals. On the other hand, the aquatic environment can also provide a platform for exchanging resistance markers between bacterial species(Organization 2014). The presence of AMR genes in water bodies has been reported globally(Herrig, Fleischmann et al. 2020). A quantitative risk assessment of AMR spreading from aquatic environment to human is in urgent need with the application of sequencing technology(Ashbolt, Amézquita et al. 2013). The hazard identification for AMR risk assessment from reservoir water in Singapore is still missing. In the genetic-based risk assessment model, investigation on both pathogenetic and non-pathogenetic bacteria is needed, as the non-pathogenetic bacteria can act as “vectors” to pass the AMR genes to human pathogens(Manaia 2017). In this perspective, the sequencing data, including the whole genome sequencing (WGS) of the resistant bacteria from the reservoir, and the shotgun metagenomic sequencing data of the reservoir microorganism community, is in urgent need to support the risk assessment system, as well as a phylogenetic-based source attribution surveillance.

As the antibiotic with the most extended history of application, beta-lactam is still one of the most common options for treating bacterial infectious diseases. Take Demark as an example; based on the “WHO report on Surveillance of Antibiotic Consumption,” penicillin contributes 61% of total antimicrobials consumption in Denmark(Organization 2018). Simultaneously, the resistance to beta-lactams is increasingly reported every year, and harboring beta-lactamases is one of the most prevalent mechanisms causing the resistance(Noval, Banoub et al. 2020). As resistant to the “Highest Priority Critically Important Antimicrobials,” 3^rd^ generation cephalosporin(Organization 2017), the Extended-Spectrum Beta-Lactamase (ESBL) and carbapenemase producers have been reported with great clinical importance.

Besides the antimicrobial susceptibility test (AST), next-generation sequencing (NGS) application can play an important role in AMR investigation. The whole-genome sequencing (WGS) on resistant strains could help track evolution history, compare the isolates from different sources, and classify them into different clades. Moreover, the DNA alignment between contigs harboring AMRs from different isolates can help to investigate the correlated AMR gene clusters. This application can benefit the surveillance of transferable resistance genes, including beta-lactamase. Compare to the mutations, the beta-lactamase is easier to spread through transferable vectors like plasmids and transposons. The horizontal gene transfer (HGT) of beta-lactamase is widely reported between different species, which brings the challenge to prevent their epidemic as the genes can be exchanged between pathogen, no-pathogen, and even some unculturable bacteria. The NGS analysis can break the boundary of species and geography and find further links between AMR gene clusters from different sources. As the sequencing technology developed, besides the WGS, metagenomic sequencing has also been widely applied in AMR studies. Besides providing the abundance level of different microorganisms, the functional gene analysis could specifically target AMR genes in complex environmental samples.

Even though NGS has been applied for AMR investigation in human and food studies in Singapore, few studies target the beta-lactam resistant bacteria from water bodies, like reservoirs, with sequencing technology. This research collects beta-lactam resistant isolates from six reservoirs and identified beta-lactamase harboring isolates with WGS. We compared their phenotype and genotype through traditional AST and sequencing technology. Furtherly, identified the specific gene cluster of beta-lactamase share between different species. Also, we have proved their transferability with the conjugation test. The multiple source comparison of sequence type and beta-lactamase genes has presented widespread beta-lactam resistance at both isolate and genetic levels. Besides, the metagenomic analysis has provided more information on the bacteria composition and AMR distribution. With our research, we aim to call national attention to control the AMR spreading through the aquatic environment in Singapore and provide more information to build up the risk assessment of AMR in the future.

## Materials and Methods

### Water sample collection

Six water samples were collected from six reservoirs respectively in Singapore during November 2018. The locations of the reservoirs were shown in supplementary. Water from 0.5 m to one meter depth to the surface, less than five meter from the bank were collected with a 1 L sterile bottle. The samples were kept at 4□ and transported to the lab within four h before counting, isolation, and DNA extraction.

### Microorganism community collection with filtration

800 ml of each water sample was filtered with 0.45 µm filter membranes to collect the microorganisms. The bacteria left on the filter membrane were dissolved in two ml saline (0.85% NaCl). The concentrated microorganism solution was further processed for colony counting and enrichment.

### Microdilution colony counting

After bacteria collection with membrane, 100 µl of concentrated microorganism solution were series diluted and drop-plated on Brilliance™ ESBL Agar (Thermo Fisher, USA) as ten µl/drop according to a 3×4 drop-plating design. For each drop with colonies between two to 20 was taken into calculation. The colonies formed on the selective were subcultured for disc diffusion.

### Culture enriched and bacteria isolation

After collected with the filter membrane, one ml of concentrated microorganism solution was added into 20 ml Mueller Hinton (MH) plus Lysed horse blood (LHB) broth (Thermo Fisher, USA) and cultured at 37□ overnight. The enriched culture was then streaked on Brilliance™ ESBL Agar (Thermo Fisher, USA) and incubated at 37□ overnight. The isolates were purified by re-streaked on MH agar and pick the single colonies for further steps. The purified isolates were subcultured in MH broth at 37□ overnight, following by mixed with 75% glycerol in a 3:1 volumetric ratio to make glycerol stock.

### Antimicrobial susceptibility test

The ESBL screening was performed with disc diffusion according to CDS methods. The amoxillin+clavulanic acid (AMC 30 µg) disc and 4 type cephalosporin discs including ceftriaxone (CRO, 30 µg), ceftazidime (CAZ, 30 µg), cefotaxime (CTX, 30 µg), and cefepime (FEP, 30 µg) were used for testing. All the discs are standard discs from Thermo Fisher, USA. Briefly, 2-3 single colonies on the overnight culture plates were picked with sterile cotton swaps and suspended in saline (0.85% NaCl) solution to reach turbidity equal to McFarland 0.5. The bacteria solution was then spread on standard MH agar homogeneously, and the antimicrobial discs mentioned before were placed on the agar surface. The agar plates were then turned over and cultured at 37 □ overnight. The diameter of the inhibit zone around discs below 9 mm was considered as the cut-off value. Isolates showed non-susceptibility to at least two antimicrobial tested were selected for sequencing and further confirmation.

The minimum inhibitory concentration tests were performed with microdilution methods and Sensititre™ ESBL Plate (Thermo Scientific, USA). Briefly, 100 µl of pure bacteria solution with turbidity equal to McFarland 0.5 was diluted with 11 ml of MH broth and mixed well. 50 µl of the diluted bacteria solution was added to each well of the MIC panel. The 96-well plates were sealed well and culture at 37 □ overnight. The lowest concentration to inhibit the visible growth was recorded as the MIC results.

### DNA extraction and sequencing

The DNA extraction was applied for both the microorganism solution and pure culture selected after the disc diffusion test. The DNA extraction followed the modified protocol published previously(Zhong 2019) with the Qiagen DNA Mini Kit (Thermo Fisher, US). The library preparation and sequencing progress were described before(Guo, Tay et al. 2019).

### Sequencing analysis of WGS

The sequencing analysis was processed with the Center for Genomic Epidemiology Server (https://cge.cbs.dtu.dk/services/). The Illumina pair-end raw reads were firstly assembled with Assembler1.2 (https://cge.cbs.dtu.dk/services/Assembler/) in Trim mode. The contigs were further processed for typing with KmerFinder 3.1 (https://cge.cbs.dtu.dk/services/KmerFinder/)(Hasman, Saputra et al. 2014), SpeciesFinder 2.0 (https://cge.cbs.dtu.dk/services/SpeciesFinder/) and MLST 2.0 server (https://cge.cbs.dtu.dk/services/MLST/) (Larsen, Cosentino et al. 2012). The genome annotation was processed by the RAST server (http://rast.theseed.org/FIG/rast.cgi) (Aziz, Bartels et al. 2008). The antimicrobial genes were detected by ResFinder (https://cge.cbs.dtu.dk/services/ResFinder/) (Zankari, Hasman et al. 2012) with filter setting as identification over 80% and mini length over 60%. The origin of replication (Ori) of plasmids was detected with PlasmidFinder (https://cge.cbs.dtu.dk/services/PlasmidFinder/) (Carattoli, Zankari et al. 2014), setting as identification over 95% and mini length over 60%. The phylogenetic tree among different species was built based on Average Nucleotide Identity (ANI) with the Genome-based distance matrix calculator (http://enve-omics.ce.gatech.edu/g-matrix/), the phylogenetic analysis within the species was processed with CSI Phylogeny 1.4 (https://cge.cbs.dtu.dk/services/CSIPhylogeny/)(Kaas, Leekitcharoenphon et al. 2014). The annotation of the phylogenetic tree was processed with iTol editor. The contigs contain AMR genes were Blast with Blastn (https://blast.ncbi.nlm.nih.gov/Blast.cgi) and Kmer to determine the location of AMR genes. The sequencing data are avabliable in the National Center for Biotechnology Information (NCBI) genome database under BioProject PRJNA719938.

### Conjugation test

The conjugation test was performed with the co-culture method, as described previously(Zhong, Guo et al. 2021). *E. coli* strain J53 was chosen as the receptor strain. Briefly, both the donor and receptor strains were picked from overnight culture plates for liquid culture in MH broth until the early log phase. Around one ml of J53 culture was added into three ml of the donor cultures, cultured at 37 □ overnight. After that, 100 µl of overnight co-culture were spread on MacConkey agar contain sodium azide (AZ, 200 mg/ml) and ceftazidime (TAZ 16 mg/ml). The successful transconjugants were confirmed with specific-primer PCR. The primers and progress of PCR were shown in Table S1.

### Metagenomic analysis

The shotgun metagenomic sequencing was performed with Illumina Miseq as described previously. The raw reads were quality checked with FastQC and processed with MG-RAST pipeline (https://www.mg-rast.org/mgmain.html?mgpage=analysis#) as the default setting. The taxa ID identified to genus level was extracted and plotted in the phylogenetic tree with (https://phylot.biobyte.de/index.cgi). The relative abundance was calculated as normalized to total bacteria reads. The alpha-diversity was presented with Shannon index, Simpson index, species richness, and 80% abundance coverage, calculated based on the species reading list from MG-RAST. The AMR genes annotation was performed with ResFinder from the CGE server.

### Bacteria community analysis

The overlap analysis of the MLST type of *E. coli* from different sources was plotted with InteractiVenn (http://www.interactivenn.net/index.html) (Heberle, Meirelles et al. 2015). Besides our study, WGS data from another five studies done in Singapore were included (Bioprject number: PRJNA476828(Mo, Seah et al. 2019), PRJNA398288(Harris, Ben Zakour et al. 2018), PRJEB26639(Guo, Tay et al. 2019), PRJEB34067, PRJNA625931(Ong, Khor et al. 2020)). Studies were grouped into three main sources as human (healthy donor and clinic patients), food (the ready-to-eat food and raw meat), and the environment (the reservoir water and wild animals). The MLST type was extracted from their publications.

To better address the beta-lactamase shared between different sources, seven studies covering six isolation sources were selected for analysis. Besides the five studies mentioned previously, two more studies as PRJNA351910 (Wyres, Wick et al. 2016)and PRJNA431029(Heng, Chen et al. 2018), were included. These two studies mainly collected the *Klebsiella* spp. isolates from the clinic in Singapore. The beta-lactamase genes information was extracted from their publication.

### Genetic environment characterization of *bla*_CTX-M_

The multi-alignment of contigs carrying *bla*_CTX-M_ was processed with Blast+ and plotted with EasyFig. The gene annotation was processed with the RAST server and Blast+ to the NCBI nucleotide database.

## Results

### The concentration of ESBL-producing bacteria

On the six counting plates, 95 isolates were picked in total. Only 11 were confirmed as beta-lactam resistant by disc diffusion and then sent for whole-genome sequencing. Seven out of 11 were detected to harbor beta-lactamase genes in the genome. However, only two isolates were found to carry ESBL genes, and the number of isolates is too small to have statistical significance. The two ESBL positive isolates were picked from the drop-plating plate of Marina Bay and Punggol, respectively. These two colonies were formed in one of the triplicate ten µl drops from the filter collection without dilution. With a brief calculation, the estimation concentration of ESBL-producing bacteria in the Marina Bay and Punggol samples was Log_10_ 1.93 CFU/ L. While the other four water samples were below the detection limit of this method. Based on the proposed standard for water quality in the Singapore reservoir provided by the PUB, the fecal coli count should below 1000 MPN/L(Alliance). Briefly, the positive rate of ESBL among bacteria possibly below 1%, but the result is not acute due to the different measurement methods. Also, the concentration we obtained of ESBL carrier from the reservoir is at least 10,000 times smaller than the reported concentration of hospital wastewater effluent (10^3^ to 10^6^ CFU/ml).

### Basic information about the isolates

Around 350 isolates were picked from ESBL selective agar. After disc diffusion, 44 isolates were sent for whole-genome sequencing (Table S1). 13/44 were Enterobacteriaceae, including *E. coli* (9), *Klebsiella* spp(3). and *Citrobacter freundii* (1), and 31/44 (70.45 %) were non-Enterobacteriaceae. The top three prevalent species were *Pseudomonas* spp. (34%), *E. coli* (20%) and *Aeromonas* spp. (14%). One of the isolates was typed to be *Bacillus pumilus* as gram-positive. All the other 43 isolates were gram-negative based on the KmerFinder result.

Among the sequenced isolates, 17 isolates were not defined as any sequence type (ST), including *Pseudomonas* spp., *Stenotrophomonas* spp., *Citrobacter freundii*, *Acinetobacter* spp. *Bacillus pumilus*, *Burkholderia cepacian*, *Chromobacterium violaceum*, and *Ochrobacterum* spp., as they haven’t been included in the CGE server database. Six of the isolates cannot be clustered as any known ST. The STs of all the *E. coli* and *Klebsiella pneumoni*a isolates have been confirmed. Two isolates of *E. coli* from two different locations (Marina and Chinese Garden) were defined as the same ST (ST1193). Meanwhile, ST1193 has been reported as an emerging human pathogen(Johnson, Elnekave et al. 2019).

The numbers of coding genes belonging to different subsystems are presented in Fig 1. Around 3000 to 7000 coding sequences were detected in the genome of isolates. The phylogenetic analysis is agreed with KmerFinder and SpeciesFinder. The isolates belonging to the same species are located on the same branch of the phylogenetic tree. Significantly, more coding genes were detected related to metabolism functions as four subsystems, including 1) Cofactors, Vitamins, Prosthetic Groups, Pigments; 2) Protein Metabolism; 3) Amino Acids and Derivatives; 4) Carbohydrates. Enterobacteriaceae were detected to have higher numbers of coding genes related to potassium metabolism, while lower numbers of genes related to stress responses compare to Non-Enterobacteriaceae.

**Fig 1.**
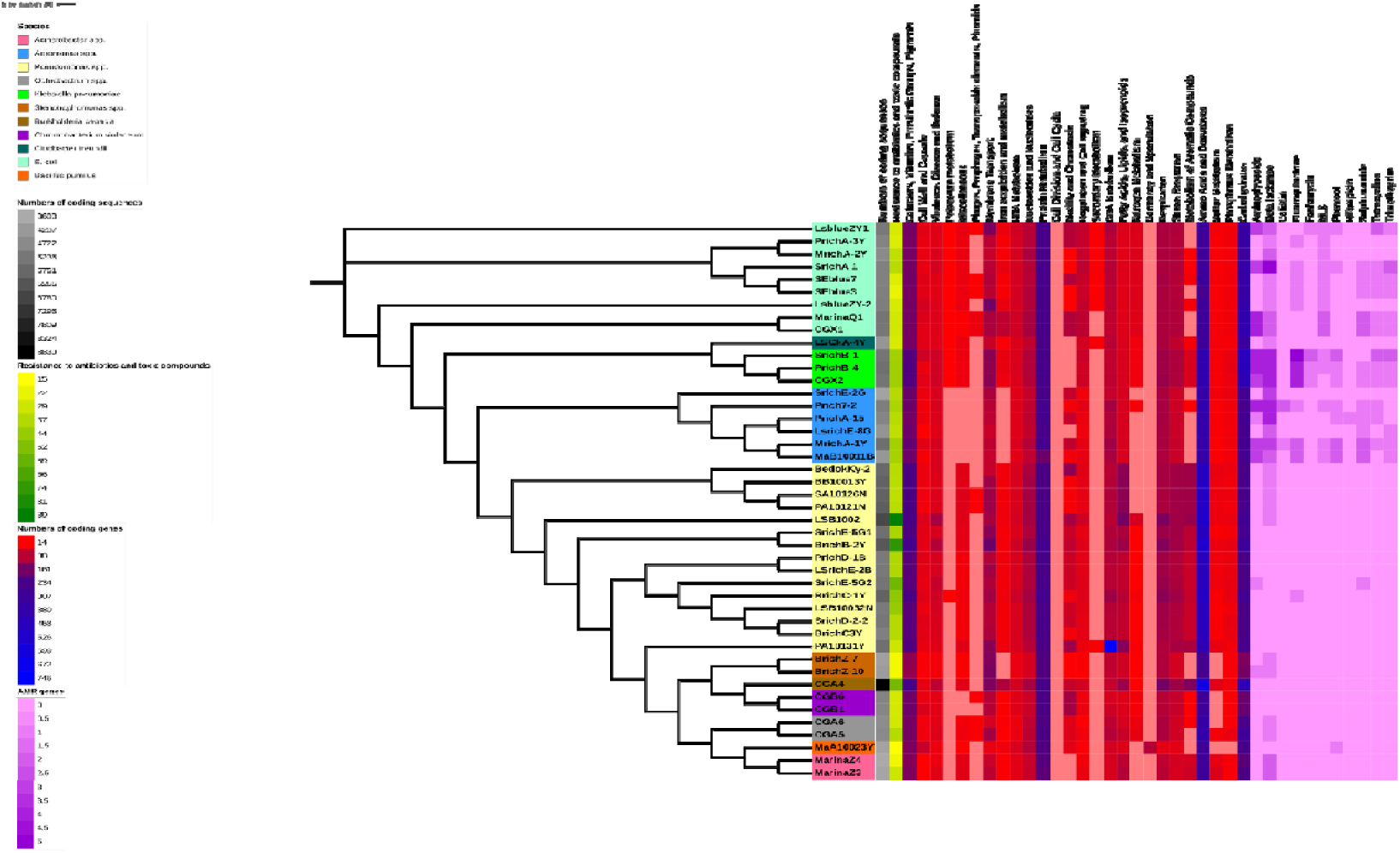
Phylogenetic analysis and coding sequences annotation. The phylogenetic tree was built based on the ANI since including all different species. The coding sequences were annotated by the RAST server grouped based on the subsystem. The AMR genes were detected with ResFinder 4.0. The color presents the numbers of coding sequences and AMR genes that belong to this function group.

Interestingly, there is no significant difference among different species for numbers of genes related to transportable elements and resistance to toxicity compounds. But in further analysis on the Ori with PlasmidFinder (Table 4), Oris was only detected in *E. coli* and *Klebsiella pneumonia* isolates. The numbers of AMR genes subjected to different classes were also plotted in the heatmap.

### The AMR genes profile of isolates

The acquired AMR genes were detected with ResFinder and are shown in Table 1. Eleven out of 44 isolates failed to show any acquired AMR genes, the 11 isolates including *Pseudomonas* spp. (8), *Chromobacterium violaceum* (2), and *Burkholderia cepacia* (1). Nine isolates were detected only to carry one resistance gene; 20 isolates were carrying AMR genes subject to more than one class. Ten isolates were harboring more than ten resistance genes. The *Klebsiella pneumoniae* strain SrichB-1 was carrying the highest number of AMR genes as 23. Colistin, fosfomycin, and rifampicin are less prevalent as detected in less than four isolates. Other classes of AMR genes were detected in more than ten isolates, and significantly, beta-lactamase genes are the most dominant AMR genes. From a genetic angle, 16 isolates are multi-drug resistant.

**Table 1.**
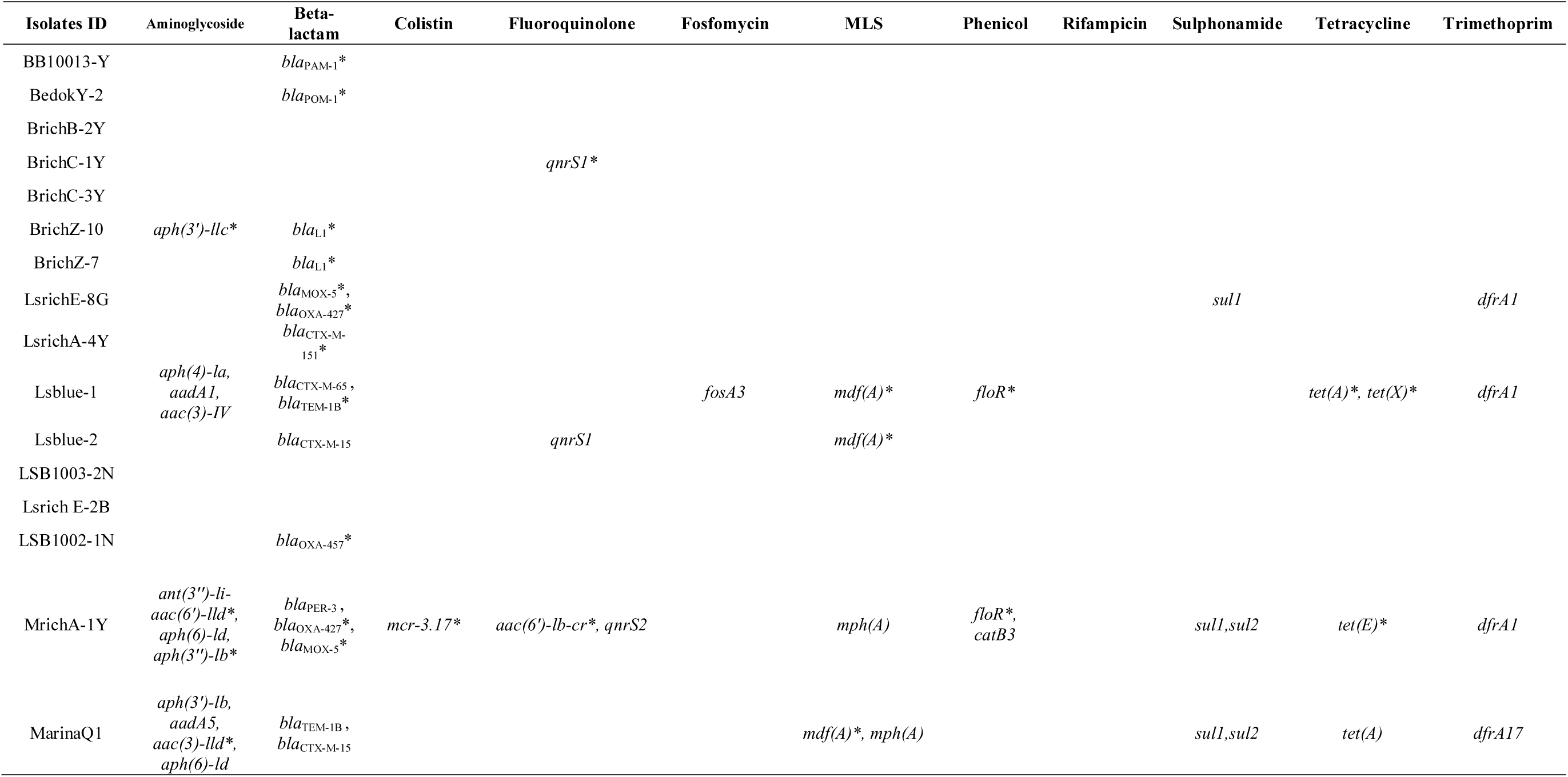
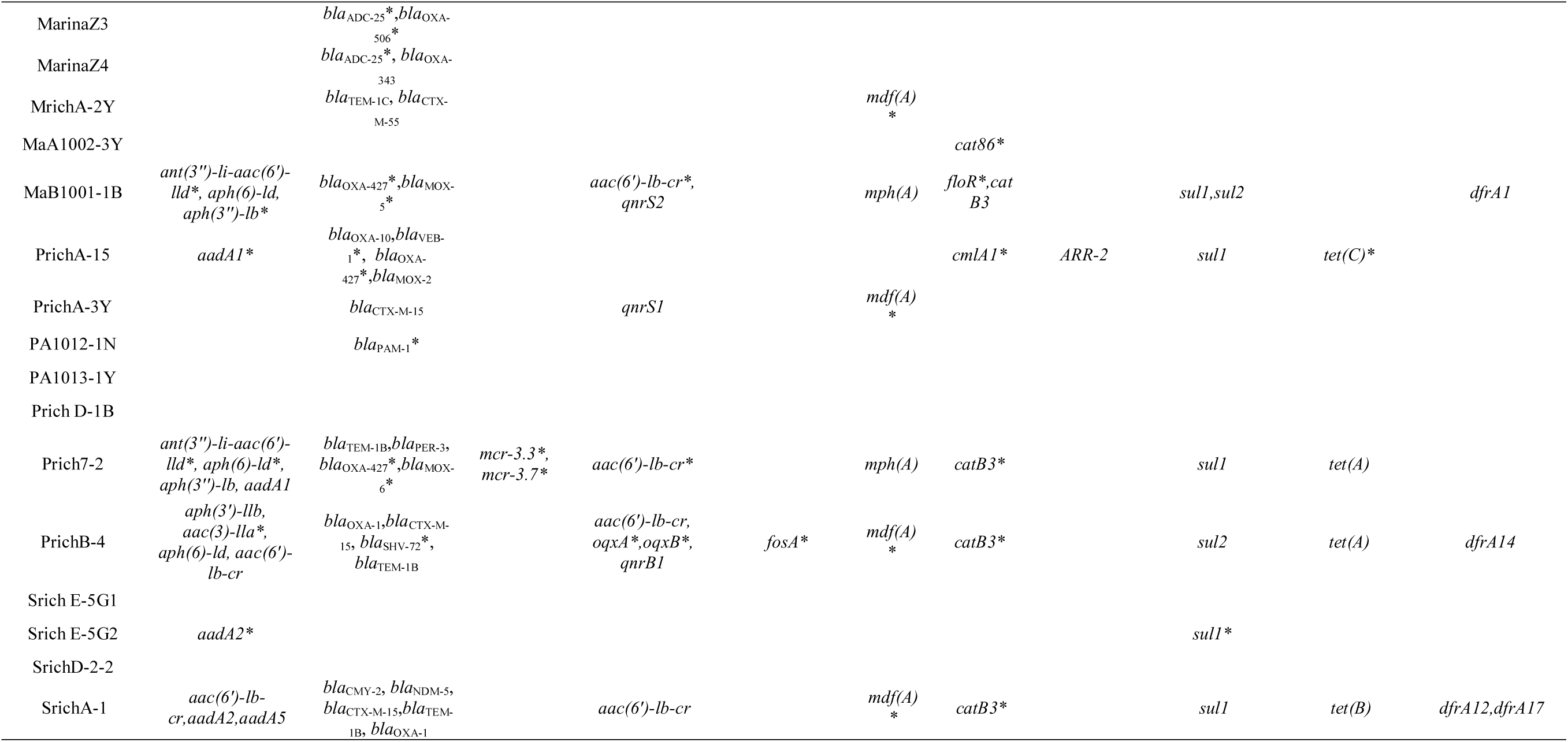
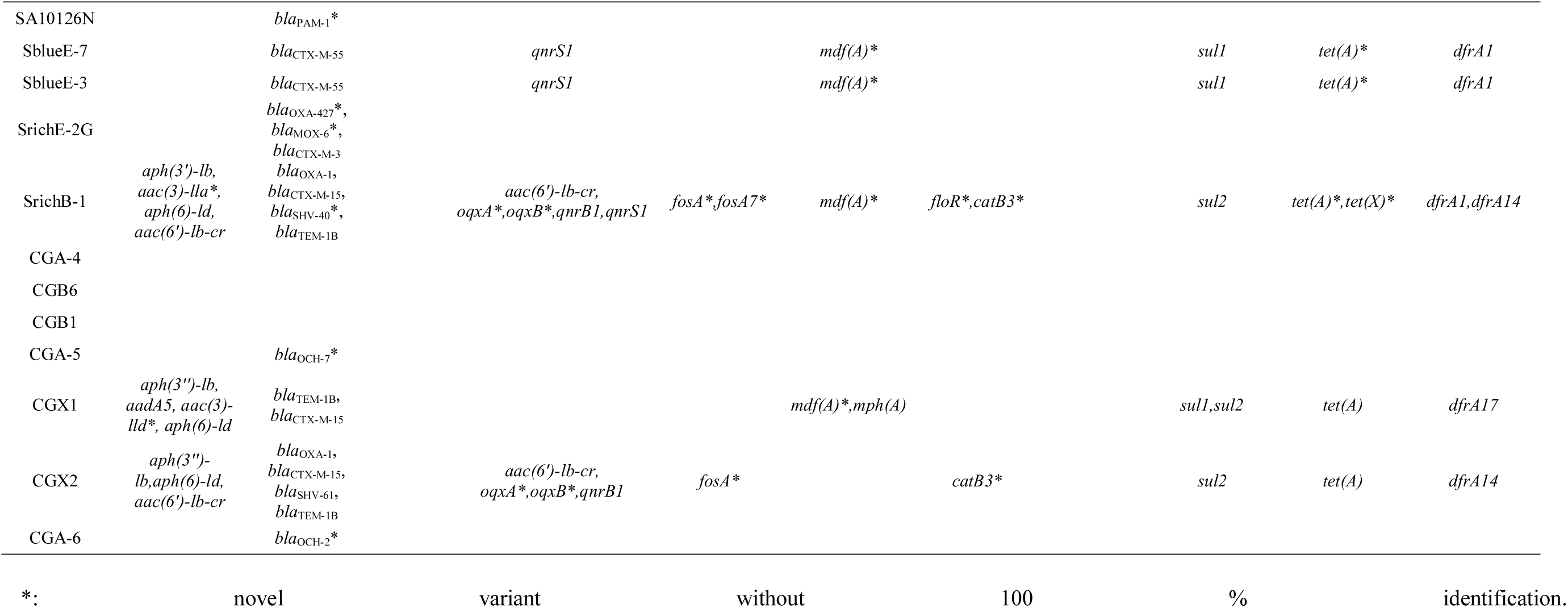
AMR genes detected by Resfinder.

Thirty out of 44 isolates were detected with known beta-lactamase genes or novel variants. The top 3 prevalent species to harbor beta-lactamase genes were *E. coli* (9/9), *Aeromonas* spp. (6/6), and *Pseudomonas* spp. (5/15). *Pseudomonas* spp. strains have shown a very high rate of disagreement between disc diffusion test and harboring of beta-lactamase genes. 13/30 isolates were detected to carry only one beta-lactamase gene, the other 17 isolates were harboring at least two beta-lactamase genes. The different classes of beta-lactamase genes were shown in Table 3. Class A is the most abundant, as 29 class A beta-lactamase genes detected in 17 isolates. Enterobacteriaceae have shown a preference for class A beta-lactamase, as all the Enterobacteriaceae isolates carry at least one class A beta-lactamase. The top 2 prevalent beta-lactamase genes are *bla*_CTX-type_ (14) and *bla*_OXA-type_(13), which have been detected in multiple species. Except for one isolate carrying class D beta-lactamase gene, all the other *Pseudomona*s spp. were harboring class B beta-lactamase, including *bla*_PAM-type_ and *bla*_POM-type_. Interestingly, *bla*_PAM-type_ and *bla*_POM-type_ were highly reported to be specific for *Pseudomonas* spp. Besides, some other beta-lactamase genes also seem specific for certain species, including *bla*_L1-type_ for *Stenotrophomonas* spp.,*bla*_ADC-type_ for *Acinetobacter* spp., and *bla*_OCH-type_ for *Ochrobactrum* spp. Some beta-lactamase genes have only been detected in one species like *bla*_MOX-type_, *bla*_PER-type,_ and *bla*_VEB-type_ genes were only detected in *Aeromonas* spp., also, *bla*_SHV-type_ genes were only detected in *Klebsiella* spp. *bla*_CTX-type_, *bla*_OXA-type_, and *bla*_TEM-type_ genes were shared among multiple species. Some variants have shown correlation to specific species like *bla*_OXA-427_-*like* genes have been detected in all *Aeromonas* spp. Isolates. Twenty-eight of the detected beta-lactamase are novel variants without 100% identification.

Two of the isolates have been detected with *mcr-3-like* genes. Both of them are *Aeromonas* spp. isolates and confirmed as ESBL-producing with both genotypic and phenotypic tests. Later MIC test of colistin has shown the isolates are still sensitive to colistin.

### Phenotypic and genotypic comparison

Thirty isolates determined to harbor beta-lactamase genes were subjected to the microdilution MIC test. The MIC results were shown in Table S1. The interpretation of resistant, intermedium, and susceptible were based on the CLSI M100 and M45 standards. Except for the Enterobacteriaceae isolates, the interpretation of susceptibility for *Stenotrophomonas* spp. and *Acinetobacter* spp. isolates were following the specific catalog for their species, while *Aeromonas* spp. is specified in CLSI M45. The susceptibility of all other isolates was interpreted based on the Non-Enterobacteriaceae catalog in CLSI M100. The comparison of phenotype and genotype is shown in Table 2. The phenotype prediction by ResFinder was also shown together with the real phenotype obtained by measures MIC for comparison.

**Table 2.**
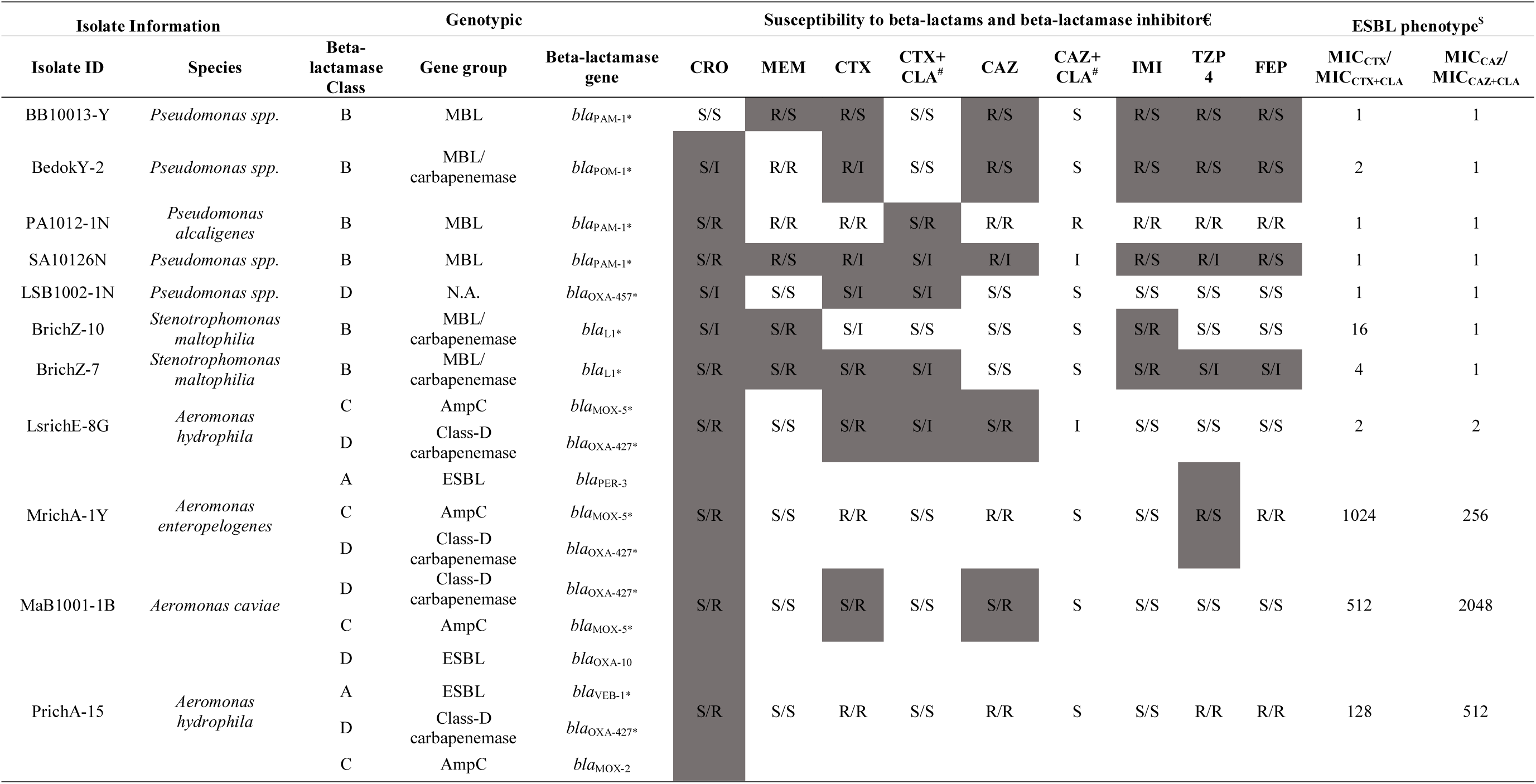
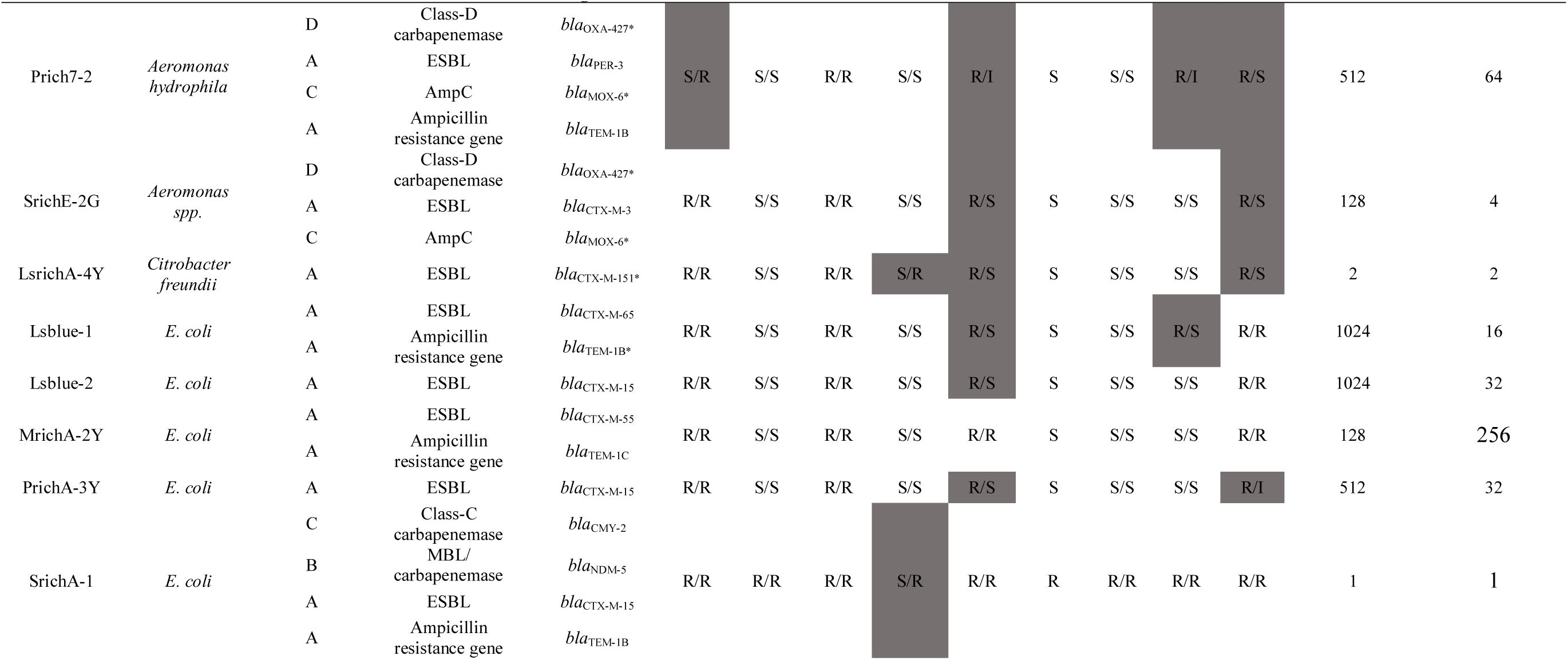
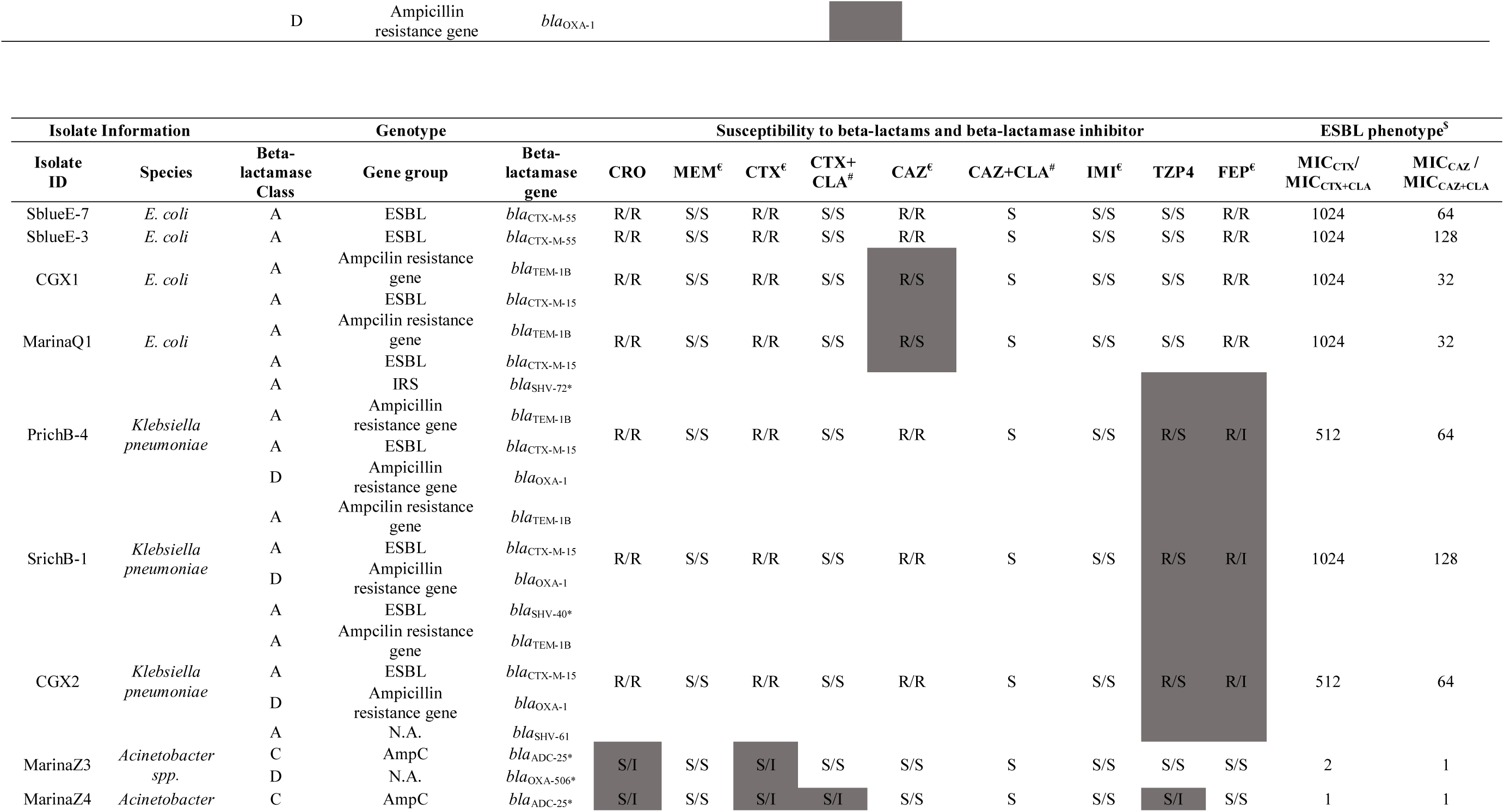
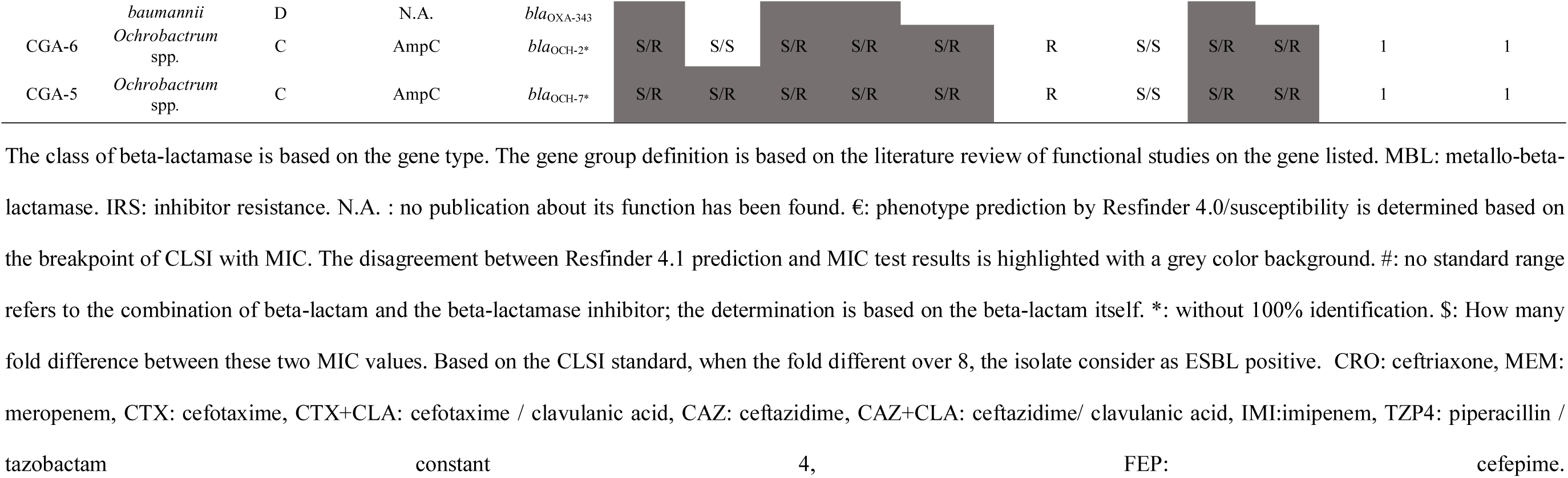
comparison of beta-lactam resistance between phenotype and genotype.

As suggested by CLSI M100, when the MIC_cefotaxime_ or MIC_ceftazidime_ is at least eight times of the MIC_cefotaxime/clavulanic acid_ or MIC_ceftazidime/clavulanic acid_, the isolate is confirmed as ESBL positive. According to this definition, 17 isolates were detected to harboring ESBL genes, but three out of 17 isolates didn’t show ESBL-producing phenotype. This may be due to the novel variant they carried, in which the mutations change the substrate and capability of beta-lactamase. Two of the isolates show the ESBL-producing phenotype without known ESBL genes. The briefly disagree rate for ESBL is around 12%. Among the ten isolates harboring potential carbapenemase, only four were finally found to be resistant to meropenem. In total, we got five carbapenem-resistant isolates; two of them were defined as *Stenotrophomonas* spp., and were harboring *bla*_L1_-*like* carbapenemase, which is B3 type Metallo-beta-lactamase (MBL). Another two are *Pseudomonas* spp. isolates with *bla*_POM-1_-*like* and *bla*_PAM-1_-like, respectively, which both belong to B3 type MBL. The other isolate is *E. coli* with *bla*_NDM-5_ carbapenemase, and this is the only carbapenem-resistance-Enterobacteriaceae (CRE) isolate we got. The *Ochrobactrum* spp. isolate with AmpC type *bla*_OCH-_ *like* genes were also shown resistant to meropenem but still sensitive to imipenem. In all, among the six sampled reservoirs, 5/6 were contaminated with ESBL-producing isolate, which has been confirmed with both phenotypic and genotypic methods.

All the isolates were harboring beta-lactamase, except for *Acinetobacter* spp. strains were resistant to at least one of the beta-lactams tested. Cephalothin, cefpodoxime, cefazolin, cefoxitin, and ampicillin were also included in tests, but there was no MIC breakpoint for non-Enterobacteriaceae in CLSI; thus, they were not shown in the comparison table of phenotype and genotype. Briefly, the MIC of all the isolates to cephalothin and ampicillin were all over the detection range (16 µg/ml). The MIC of all the isolates except *Acinetobacter* spp. were all over the detection range as 32 µg/ml. The MIC of all the isolates except MarinaQ1(*E. coli*) were all over the detection range as 16 µg/ml. For cefoxitin, all the Enterobacteriaceae isolates except strain LsrichA-4Y were sensitive to it. All the *Aeromonas* spp. isolates except one were also susceptible to cefoxitin. For the other non-Enterobacteriaceae, the MICs were over 64 µg/mL.

All the isolates containing *bla*_CTX-M-type_ genes were resistant to both ceftriaxone and cefotaxime, regardless of species. All the *Klebsiella* spp. isolates were resistant to ceftazidime as harboring *bla*_CTX-M-15_ and *bla*_SHV-type_ genes. At the same time, three *Aeromonas* spp. isolates (LsrichE-8G, MrichA-1Y, and MaB1001-1) contain both *bla*_OXA-427_*-like* and *bla*_MOX-5_-*like* genes were also resistant to ceftazidime, two *Aeromonas* spp. isolates (Prich7-2 and SrichE-2G) contained *bla*_MOX-6_*-like* instead and were less resistant to ceftazidime, suggesting that the *bla*_MOX-5_*-like* gene is important for the resistance to ceftazidime. Interestingly, the MrichA-1Y strain, which contains the combination of *bla*_OXA-427_*-like* and *bla*_MOX-5_-*like* genes together with *bla*_PER-3,_ has shown resistance to cefepime. However, when changed the *bla*_MOX-5_-*like* to *bla*_MOX-_6-*like* in the combination, as strain Prich7-2, the isolate becomes sensitive to cefepime. All the other *E. coli* isolates with the combination of bla_CTX-M_ genes, and bla_TEM_ genes have shown resistance to cefepime. However, this combination seems to be less effective on cefepime in *Klebsiella* spp. isolates as their MIC only reach the intermedium level. Except for the *Ochrobactrum* spp. isolates and SrichA-1 strain with *bla*_NDM-5_, all other isolates are sensitive to the combination of piperacillin and tazobactam constant(4 µg/ml). The SrichA-1 strain has shown resistance to all the antimicrobials tested, besides, due to the carbapenemase, the ESBL-producing phenotype has been covered up. The *Ochrobactrum* spp. isolates were resistant to all the antimicrobials tested except imipenem, this may attribute to the novel variants of the *bla*_OCH-type_ gene.

The strains have shown different responses to beta-lactamase inhibitor, clavulanic acid. All ESBL-producing isolates have been strongly inhibited of the resistance to cefotaxime by clavulanic acid, as the MIC decreased 512 or 1024 times when cefotaxime combined with clavulanic acid, except SrichE-2G strain, still decreased 128 times. The MICs of ceftazidime were less influenced by inhibitor as decreased no more than 256 times, except for the MaB1001-1B strain, which has a MIC decreased by 2048 times, even over the decrease of cefotaxime. Interestingly, the MaB1001-1B were failed to detect with known ESBL-genes. Besides, the strain LsrichE-8G, who carrying the same type of beta-lactamase as MaB1001-1B has shown a slight response to the inhibitor. For isolate LsrichA-4Y, which harboring a variant of *bla*_CTX-M-151_ showed a very slight response to the clavulanic acid.

### Phylogenetic analysis of *E. coli* isolates with AST result from multiple sources

To better describe the relatedness of *E.coli* from different sources, WGS data of isolates with AST result from PRJNA39288, PRJEB26639 was extracted and compared with isolates from reservoir water based on core-genome SNP. The reservoir isolates included current research and the isolates reported by our group previously(Zhong, Guo et al. 2021). As in Fig 2A, in total, 95 isolates were included in the phylogenetic analysis, as 18 isolates from the reservoir, 23 isolates from the raw meat, and 44 isolates from the clinical patients. Based on the phylogenetic tree, there is no clear border between different sources. Based on the susceptibility interpretation, all except three clinical isolates were resistant to CTX. However, two of the three sensitive isolates except Mer-99 were harboring the beta-lactamase gene, as OXA-type combine CTX-M-15 and CTX-M-27, respectively. Meanwhile, all the other isolates carrying the same type of genes were resistant to CTX. This study has no explanation as to why these genes are not functional in the three isolates. Around 96% of the 95 isolates were susceptible to TZP, except four isolates from the reservoir shown resistance to TZP. However, there is no clear correlation between the beta-lactamase gene they are carrying and their resistance to TZP, as their beta-lactamase genes are diverse. To better understand the relationship between reservoir isolates and human source isolates, a phylogenetic analysis on isolates from both reservoirs and healthy donors was performed (Fig S1). The same STs were found to be shared between different sources. The detailed comparison of ST type from multiple sources has been shown in Fig 2B.

**Fig 2.**
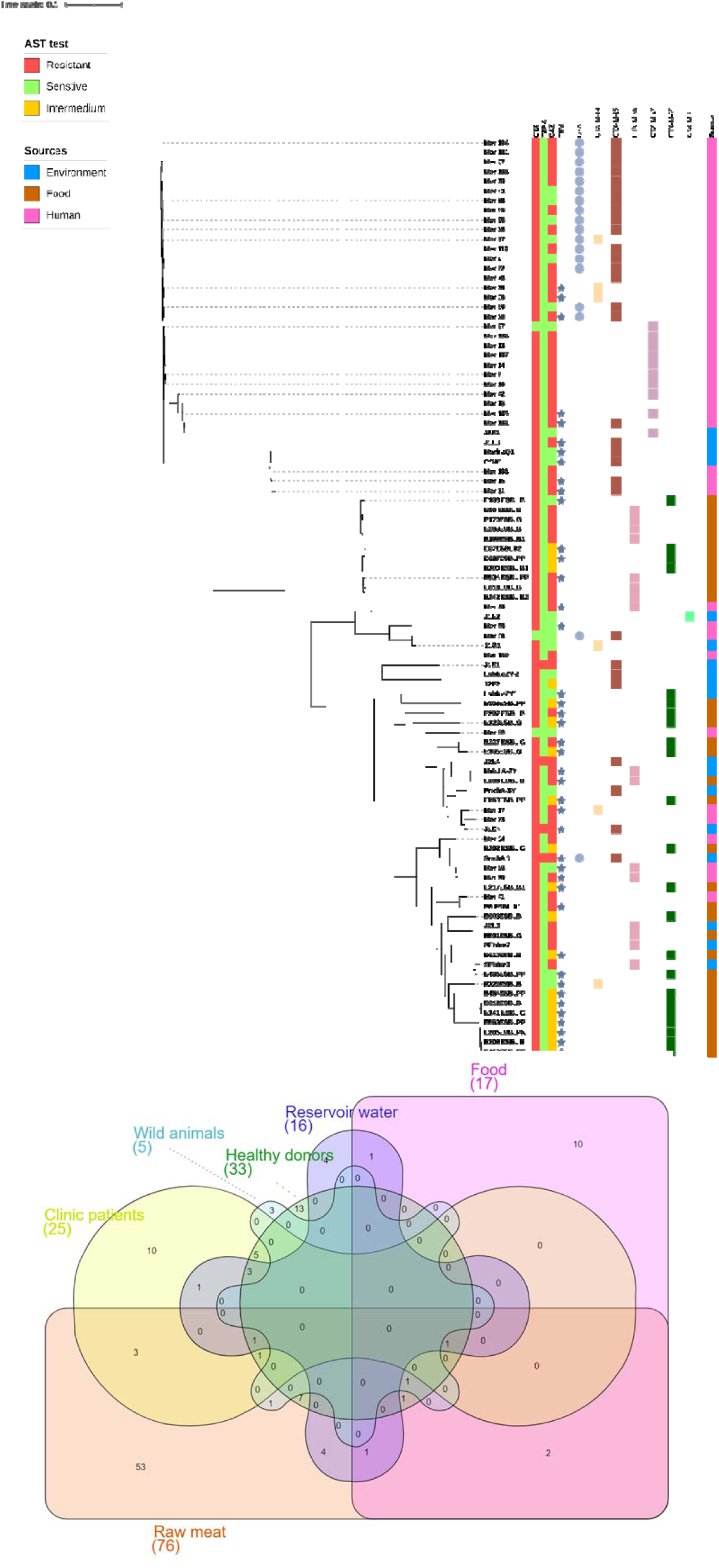
**A.** Phylogenetic tree of *E.coli* from different sources annotated with AST results and beta-lactamase genes. The *E. coli* isolates from human, food, and water sources were aligned based on core-genome SNP. Only the isolates reported with AST interpretation were extracted for the phylogenetic analysis. The MG1655 strain is used as the reference. **B.** Overlap analysis of MLST type of E. coli from different sources. The ST types of E.coli isolates from multiple sources, including human (healthy donors, clinic patients), ready-to-eat food, raw meat, wild animals, and reservoir water, were extracted for the analysis. The total numbers of ST types are shown in the bracket next to the subgroup label. The numbers of ST shared between more than one source are labeled in the overlap area.

To further investigate the potential transferring of *E. coli* among different sources, we briefly compared the ST type of *E. coli* collected in Singapore from various sources, including food, humans, and the environment. Each source contains two subgroups: food: ready to eat (RTE) food and raw meat, human: healthy donors and clinic patients, and environment: reservoir water and wild animals. The involved Bioproject details have been described in the materials and methods. Three ST types have been found to distribute among humans, food, and animals, including ST10, ST38, and ST23, especially ST10 has been found in 5/6 subgroups that exclude the wild animals. Besides these three ST, human and food sources shared other nine ST as ST93, ST48, ST101, ST117, ST349, ST394, ST354, ST2179, ST2732, and ST3580. While except the three ST share by all sources, the numbers of shared ST between human and the environment, food, and the environment was smaller, as four and five respectively. As detailed, ST1011, ST127, ST155, ST1730, ST206 only shared between food and the environment, while ST131, ST410, ST648, ST1193 only shared between the humans and the environment. This may due to the larger sample size of human and food isolates, but it may also raise the question of whether the bacteria from environmental sources are more difficult to colonize on humans. However, if the percentage of ST for overlap is compared, 75% of the ST found in reservoir water can be found in other sources as both humans and food. Comparatively, only 41% of the ST found in ready-to-eat food is shared with other sources. 37.5% of the ST from reservoir water can be found in either clinic patients or healthy donors, while 19.7% of the ST from raw meat and 17.6% of the ST from RTE food, respectively. This is different from our hypothesis; traditionally, RTE food should have more exportation on humans. The sample size of wild animals is the smallest as only five ST has been reported, but contain an ST23 shared among four subgroups exclude reservoir water and clinic patients. Except the three ST only belong to the wild animal group, the other ST, like ST602 from the wild animals, can be found in the raw meat group. On the other hand, the raw meat and the wild animal are the only two groups containing ST602. Due to the different sample sizes, it is difficult to evaluate which source contributes more to the E. coli spread to humans based on current data. Interestingly, within the subgroup, the RTE food and raw meat only shared six ST. Relatively, the raw meat group shares seven ST with the reservoir water; considering the large sample size of raw meat, the isolates type between the cooked and raw materials could be quite different. Even though many limitations for the overlap analysis, we can still find the wide distribution of specific types of *E.coli* like ST10 regardless of sources and has caused human infections in the clinic.

### Phylogenetic analysis of *Klebsiella pneumoniae* from humans and reservoir water

To better indicate the relationship between *Klebsiella* spp. from different sources, a phylogenetic analysis based on core genome SNP was performed on our isolates from the reservoir and other isolates from the clinic (Fig S2). As one of the top ESBL-producing pathogens in the clinic, the sequencing record of *Klebsiella pneumoniae*. we extracted can be up to the year 2011 to 2012. Unlike the *E .coli* strains, no same ST has been found between humans and reservoir water. Interestingly, except for isolate SRR7371314, neither the same ST was found between healthy donors and patients. It seems like the isolation spreading bound existing not just between the environment and humans, but also between individuals with different health conditions. But the same combination of beta-lactamase genes has appeared in various sources. For example, the combination of *bla*_OXA-1_, *bla*_TEM-1B,_ and *bla*_CTX-M-15_ has presented in 14/95 isolates in the analysis, regardless of sources. This raises the hypothesis that *Klebsiella pneumoniae* is acting more like a “vector” of passing AMR genes instead of directly colonizing humans. This hypothesis has been further supported by beta-lactamase distribution analysis and the conjugation tests. The *bla*_SHV-like_ genes were not plotted in the phylogenetic tree as all of the isolates are carrying *bla*_SHV,_ and most of them are novel variants. Besides, the *bla*_SHV_ of our *Klebsiella pnerumoniae* strain are all determined to be located on the chromosome, and the variant type has shown ST specification in both human and reservoir isolates, which make it more like an intrinsic resistance gene, the rich mutations of the *bla*_SHV_ may due to the evolution of the genome.

Quite a pity that we didn’t find many sequencing data reports of *Klebsiella* spp. from other environmental sources and food in Singapore, which stop us from tracking their transportation among different sources.

### Ori detection and location determination of beta-lactamase

The beta-lactamase genes’ locations were determined by the best hits (Table S4) found by NCBI Blastn, also with the Ori detected by PlasmidFinder (Table S3) as reference. Briefly, based on the best hits found by Blastn, the Enterobacteriaceae are more likely to carry transferable beta-lactamase located on the plasmid. The Non-Enterobacteriaceae are mainly detected with beta-lactamase located on the chromosome. This is also agreed with the numbers of Ori detected in different groups. Known from the Blastn result, the *Pseudomona*s spp., *Stenotrophomonas* spp. and *Ochrobactrum* spp. isolates each has a chromosome-located beta-lactamase. As mentioned previously, these beta-lactamases are mainly specific to the species. Each of the two *Acinetobacter* spp. isolates were harboring new variants of *bla*_OXA-type_ and *bla*_ADC-type_ genes, co-located on the chromosome.

Except for *bla*_TEM-1B_ detected in Prich7-2 strain were similar to a plasmid reported, all other beta-lactamases were probably located on the chromosomes. This is not exactly in agreement with the PlasmidFinder results as no Ori has been detected in *Aeromonas* spp. isolates. Interestingly, the *bla*_PER-3_ genes detected in Prich7-2 and MrichA-1Y strain were likely from an integron sequence, a highly similar integron has been reported in *Aeromonas hydrophila* strain RJ604. The integron sequence is a very specific transferable cluster, which could acquire AMR genes and insert them into the genome.

Three out of five *bla*_CTX-M-15_ genes in *E. coli* isolates were determined to be located on the chromosome. The phenomena have been found and addressed by our previous research(Zhong, Guo et al. 2021). The *bla*_CTX-M-15_ genes were inserted into the chromosome driven by a specific insert sequence. The structure and genetic environment were further compared then (Fig 6). All the other beta-lactamase except *bla*_CMY-1_ were found highly similar to the sequences of plasmids from multiple species and sources. Also, except for Sblue-3, all the *E. coli* isolates have been detected to have at least one Ori. This may suggest their rich capability for genetic communication. Even though the contigs carrying *bla*_CTX-M-55_ of Sblue3 were found to be similar to plasmid pCFSA1096 from *Salmonella enterica* strain CFSA1096, as no Ori was detected in the Sblue-3 strain, this may support our conclusion from the previous research (Chapter II), that this gene cluster may initially located on a plasmid and then jump to the chromosome during HGT.

Different from *E. coli,* the *bla*_CTX-M-15_ genes in *Klebsiella* spp. isolates were all co-located with *bla*_TEM-1B_ on the contigs similar to the sequence of the plasmid, also the contigs from different strains are highly similar to each other, which may suggest they are transferred from the same ancestor. This widely distributed plasmid has also been shown to be conjugatable then.

### Conjugation capability test

The *Klebsiella pneumoniae* isolates have shown the capability of passing ESBL genes to a different species. The genes have been conjugated *to E. coli* strain J53 successfully in a frequency around 10^-6^. The successful transconjugants *E. coli* can survive on agar plates containing 16 µg/ml of ceftazidime, a further PCR and MIC test has also found the ESBL phenotype and resistance to cephalosporin, which may reflect the donor has obtained the resistance and the beta-lactamase can be successfully activated in a different species.

### Transferable beta-lactamase is distributed with different prevalence in different sources

From the conjugation tests, we proved conversed ESBL gene clusters to be conjugated among different species. Based on this conclusion, we compare the beta-lactamase shared between isolates from different species, including *E .coli* and *Klebsiella* spp., we also include the beta-lactamase found in the *Aeromonas* spp. in this study (Fig 3). However, there are pretty limit WGS report of *Aeromonas* spp. from Singapore, so we failed to find sequencing information for comparison. Besides, we also noticed that the beta-lactamase carried by our *Aeromonas* spp. isolates can be found in *Klebsiella* spp. isolates from the clinic, this is also one reseason we included *Aeromonas* spp. in this analysis. Due to each study’s sample size being different, we use the positive rate of the gene carrier to plotted the gene communication and distribution among various sources.

**Fig 3.**
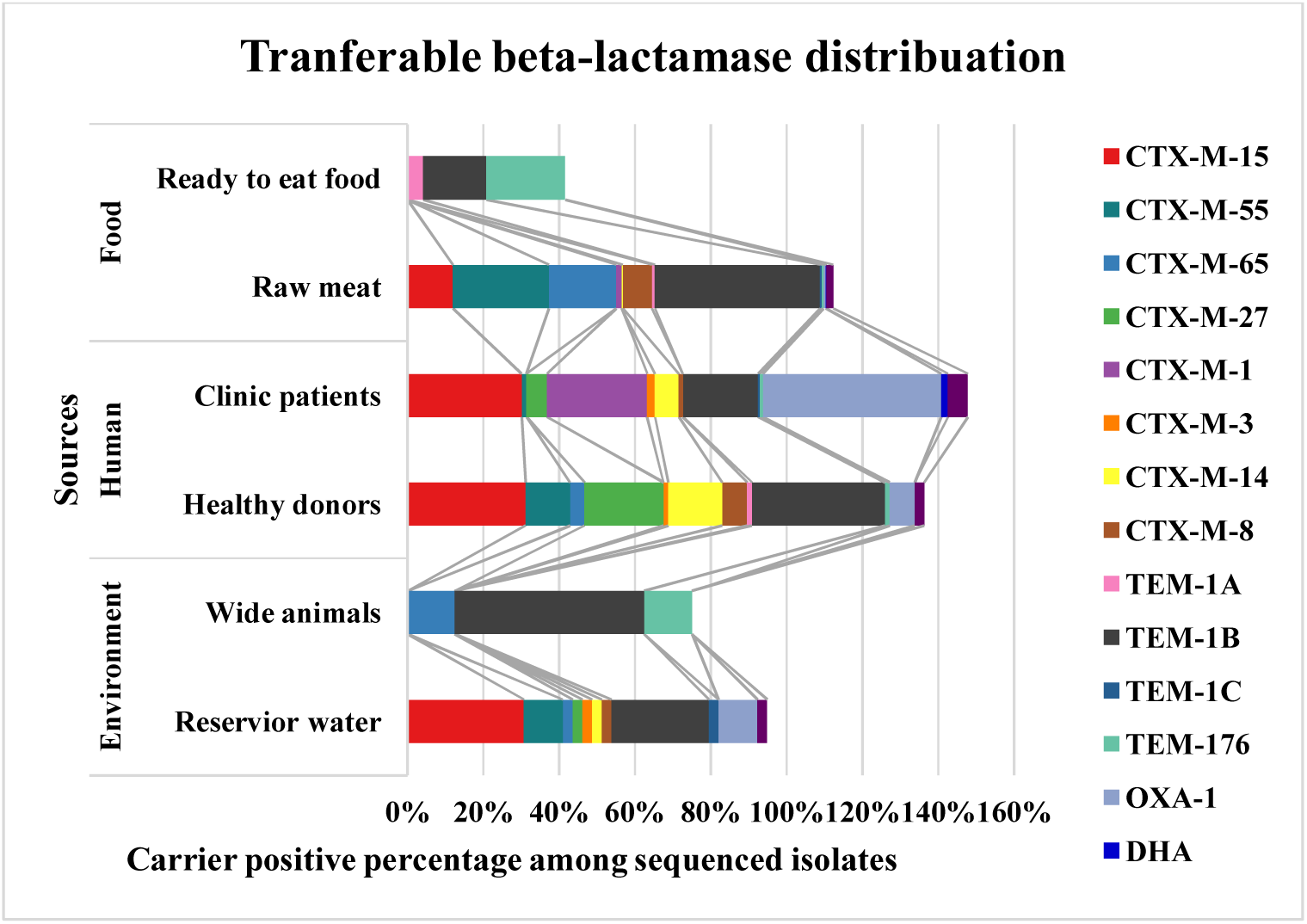
The beta-lactamase carrier present in isolates from different sources. The beta-lactamase genes information of isolates from this study and other studies was collected, including *E .coli*, *Klebsiella* spp. and *Aeromonas* spp (only from this study). Due to the difference in sample size, the positive rate of carrier among total sequenced isolates was used for comparison. The same beta-lactamase gene was linked with line and labeled with the same color. The beta-lactamase detected only in one group was not present here. The total percentage is over 100% is due to the co-existing of genes.

Exclude the wild animals and RTE food, the other four subgroups have been found to contain a rich type of beta-lactamase. As the most come beta-lactamase gene, *bla*_TEM-1B_ has been found in all six subgroups, including all three species. Besides, most of the beta-lactamase was shared between all three sources, exclude *bla*_CTX-M-27_ and *bla*_CTX-M-3_ are only shared between the environment and humans, *bla*_CTX-M-1_, *bla*_TEM-1A_, and *bla*_DHA_, only shared between food and human. This result reflects there is no bound for the beta-lactamase spreading among different sources and species.

### Genetic environment analysis of the *bla*_CTX-M_ gene

As the capability of passing ESBL genes between different species was shown, a DNA alignment of contigs harboring *bla*_CTX_-_M_ genes was also investigated. A conserved gene cluster containing the same types of genes has been detected in *Aeromonas* spp. *E. coli* and *Klebsiella* spp. Including the ISEcp1 insert sequence, a *bla*_CTX-M_ gene, and a *Wubc-like* gene. Interestingly, the sequence of this cluster found on the plasmid of *Klebsiella* spp. is the same as the one located on the chromosome of *E. coli* isolates, which possibly suggests this gene cluster could be moved from the plasmid to the chromosome. The same gene cluster has been detected in eight (36%) isolates among all the isolates harboring *bla*_CTX-M_ genes collected in this and previous research of reservoir bacteria, suggesting the wide local spreading of this gene cluster.

A further blast has been done between isolates from different sources and species (Fig 4). The genetic environment of CTX-M-15 and CTX-M-55 was found to be highly similar and conserved. They were both highly evolved with the *ISEcp1* and followed by a *WbuC* gene. And this gene cluster has been widely reported, including in our previous research(Zhong, Guo et al. 2021). The proximity of CTX-M-15 or CTX-M-55 is widely presented among different species and sources. The genetic environment of CTX-M-65 has also been shown conserved among different species but highly different from CTX-M-15 and CTX-M-55 (Fig S2).

**Fig 4.**
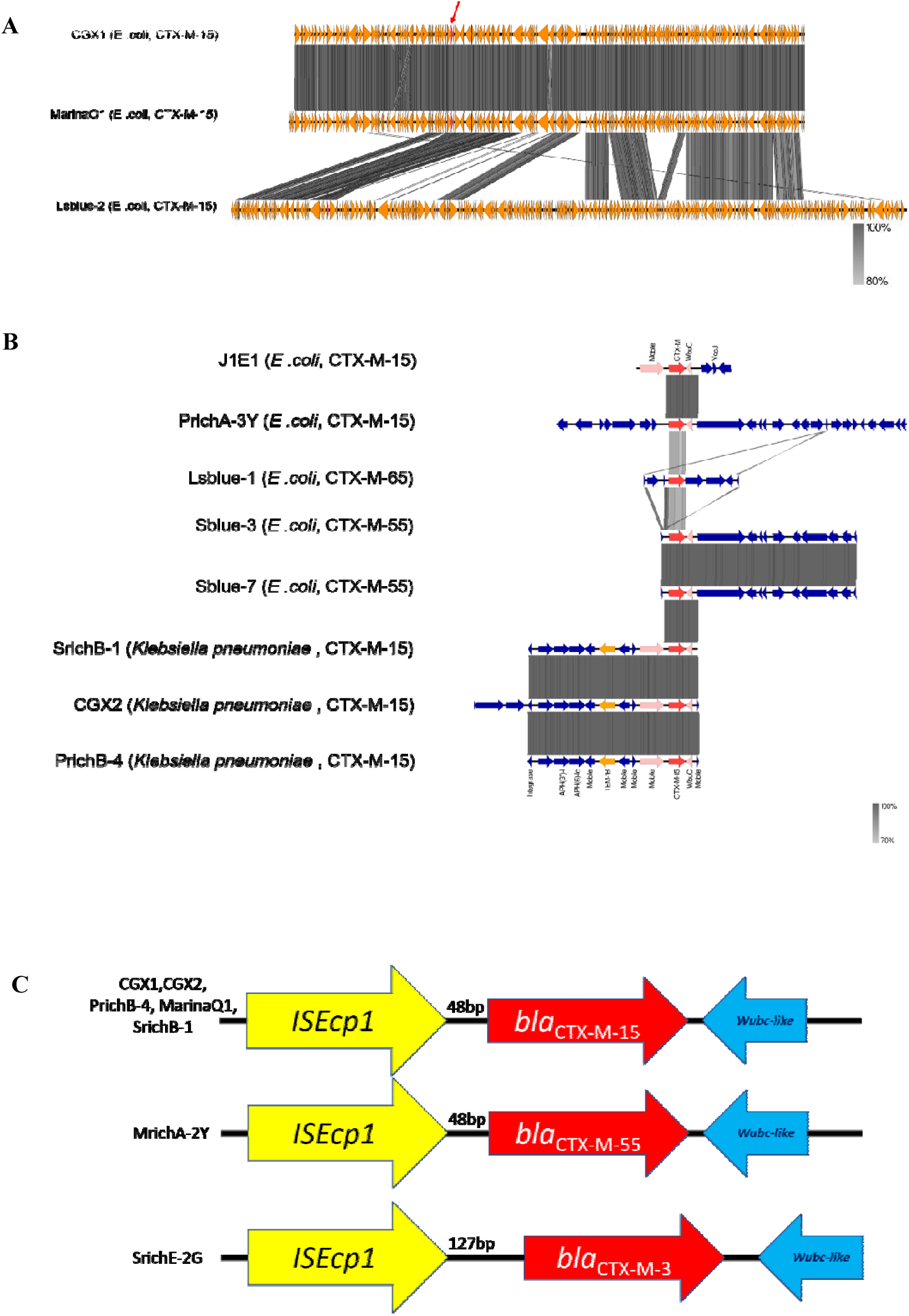
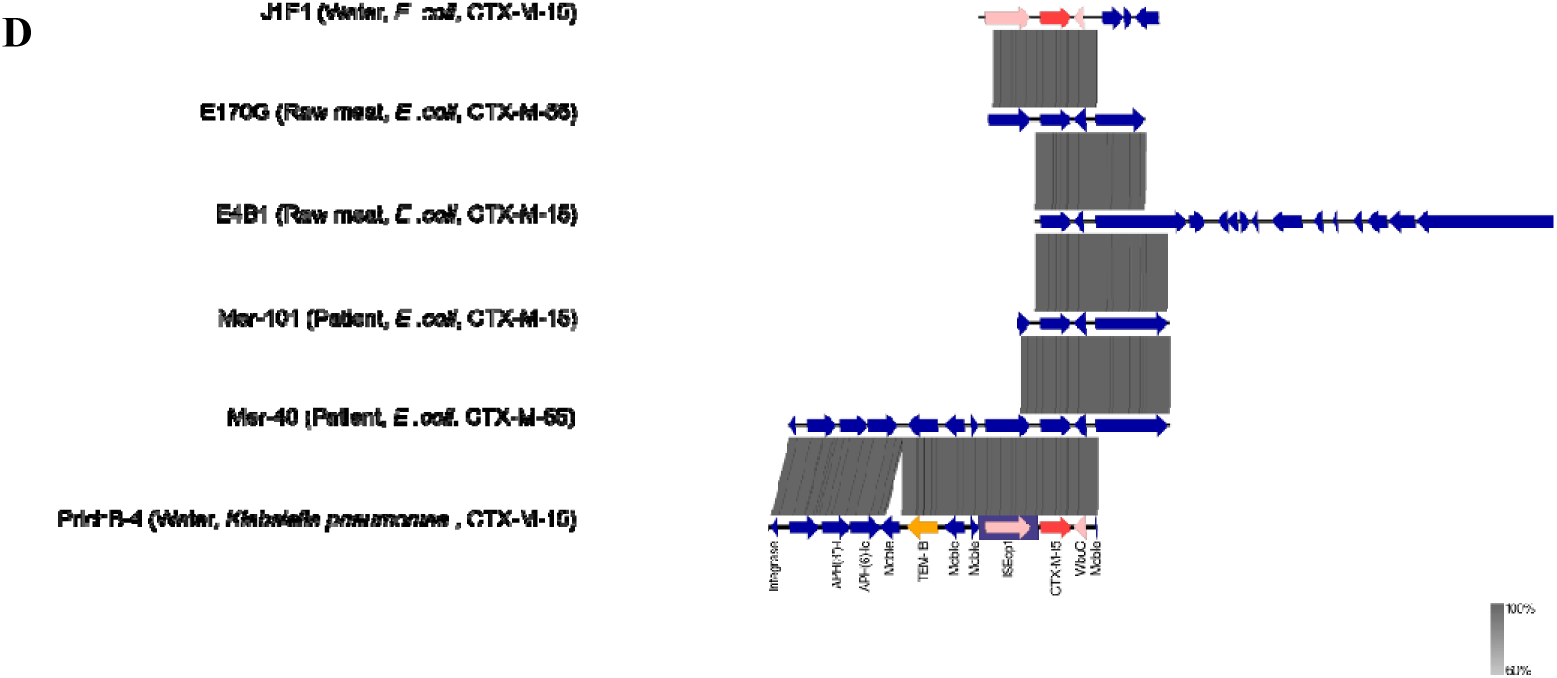
DNA alignment of contigs with *bla*_CTX-M_ genes from different sources’ isolates. The contigs contain CTX from different isolates were extracted for DNA alignment with Blast+ and plotted with EasyFig. **A**: Alignment of contigs over 20 kbp contain CTX-M-15 from reservoir isolates of *E .coli* in this study. **B**: Alignment of contigs contain CTX-M-15, CTX-M-55, and CTX-M-65 between 3 kbp to 12 kbp from reservoir isolates of both *E .coli* and *Klebsiella pneumonia*. **C**: The genetic elements of the transferable gene cluster. The conversed gene cluster from multiple species was extracted to present with gene annotation and distance between genetic elements. **D:** Alignment of contigs contain CTX-M-15 and CTX-M-55 from multiple sources and species.

### Taxonomy analysis of shotgun metagenomic data of reservoir samples

Based on the MG-RAST taxonomy analysis result, around 1500 species taxa unit belong to 590 genera has been identified (Fig 5). The total reading number is between 2588981 to 5000134 was called. Proteobacteria are the dominant group. Besides, Firmicutes, Actinobacteria, and Bacteroides are also detected with rich abundance. The Shannon index, Simpson index, species richness, and 80% abundance coverage was calculated based on the taxa unit identified to species level. The community of the Serangoon reservoir has shown unexpected low diversity due to dominated by *Microcystis aeruginosa*, which has covered 20% of the relative abundance; this reminds us of the light green color of the sample from Serangoon. Also, the species richness of Serangoon is the highest, which presents as a significantly lower score of Shannon, Simpson and, 80% abundance coverage.

**Fig 5.**
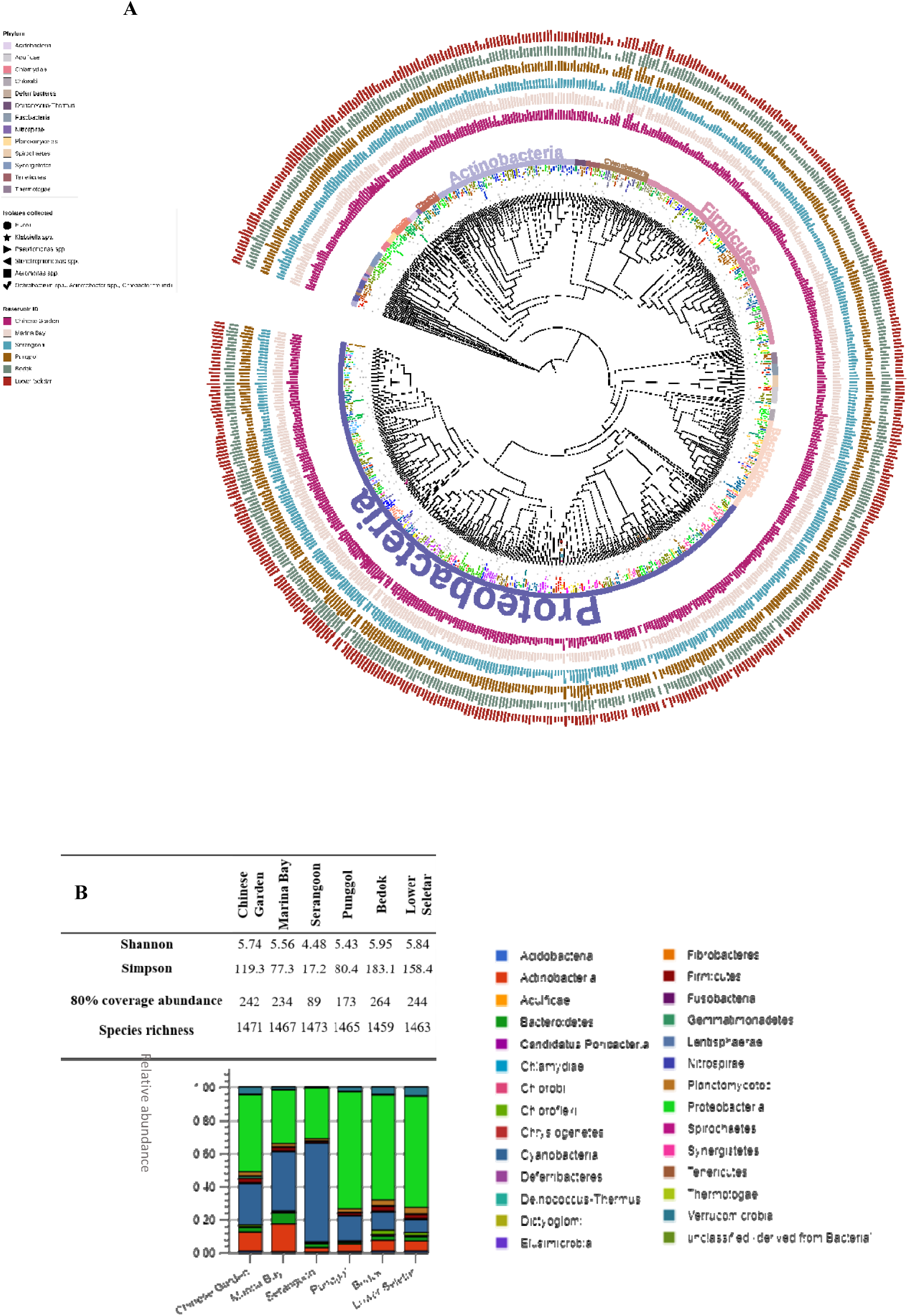
Phylogenetic analysis of taxa ID identified to genus level with shotgun metagenomic sequencing analysis on different reservoirs, annotated with relative abundance normalized to total bacteria. **A.** The phylogenetic tree was build based on the taxa unit identified with MG-RAST at the genus level; 584 leaves are included. The inner-circle presents the phylum of the taxa unit, phylum with over five leaves were annotated directly on the ring. Phylum with fewer leaves was color labeled as the legend. The six outer-circle presents the Log_10_ number of taxa units in six reservoirs, from inside to outside, marked with the color legend. The isolates from each reservoir were also labeled on the brach with the same color of their reservoir source. B. Alpha-diversity analysis of the bacteria composition. The composition of bacteria of the water samples was annotated with the MG-RAST pipeline, and composition at phylum level was plotted. The alpha-diversity indexes were calculated based on the readings identified to species level. A significantly lower diversity level was found in the sample of Serangoon.

### Functional gene annotation of shotgun metagenomic data of reservoir water

Between 1106317 to 1930613 functional unit hits were detected based on the SEED database (Fig S4). Like the WGS data, the dominant group is related to the essential metabolism of nutrients, like carbohydrate metabolism. We specifically look into the function group of antimicrobial resistance and found the beta-lactamase can be detected in all six samples with different relative abundance. The multi-drug resistance efflux is still the most common group among the functional hits belonging to the “ Drug-resistant” group (Fig S5).

A further annotation targeting the AMR genes with ResFinder is shown in Table 3. Interestingly, the AMR genes detected are entirely different from the ones seen in isolates. Bedok and Lower Seletar samples failed to detect AMR genes with assembled contigs. Even though the sample was not applied with enrichment and selection, the beta-lactamase genes were still prevalent. However, the types of the gene are different from the genes found in the isolates. The *oqxA*-like gene was detected in some isolates from Punggol and Serangoon. Significantly, the raw reads with KMA methods can help to detect more AMR genes but lack genetic information. Besides, the assessable contigs carrying AMR genes are too short to determine their source organism.

**Table 3.**
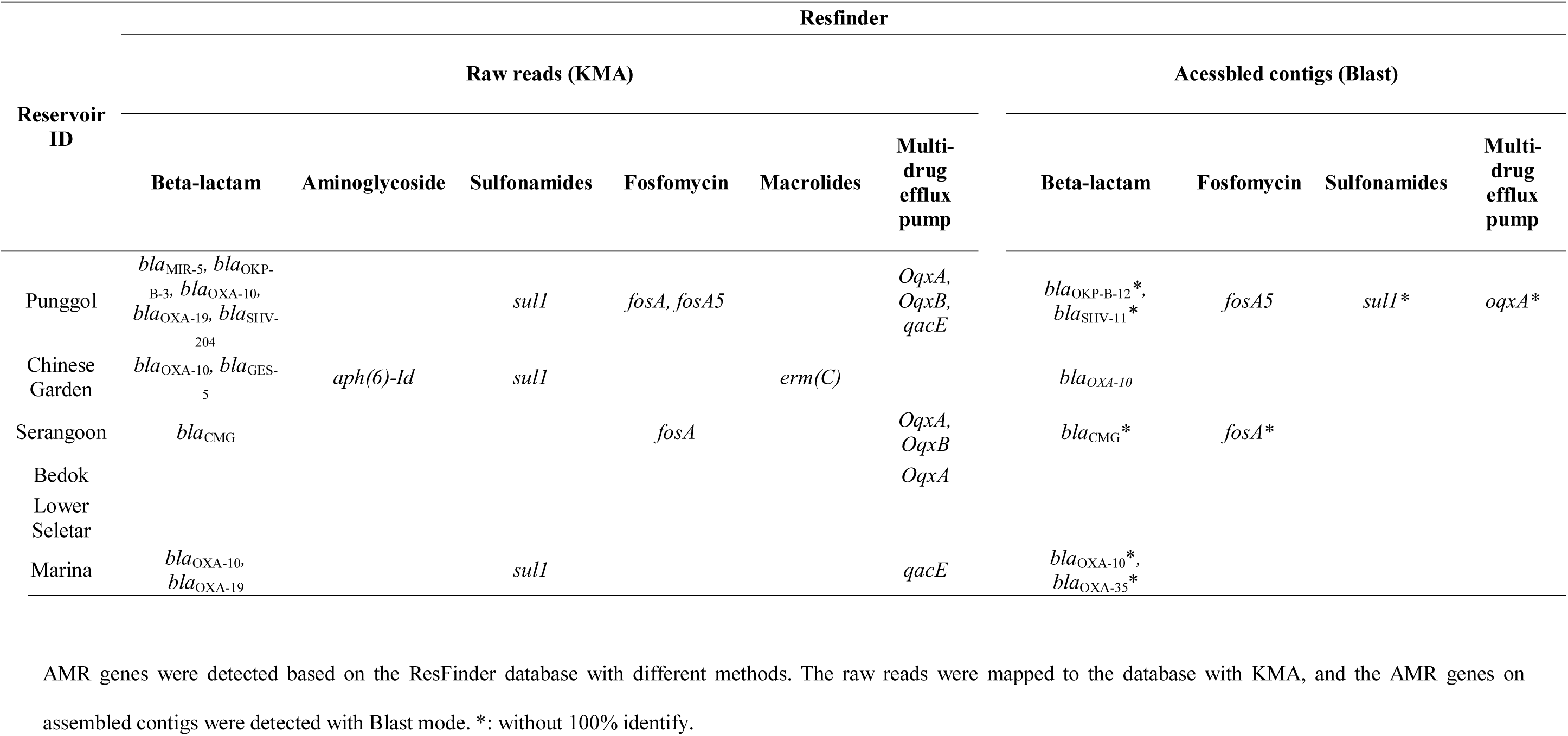
AMR gene detected in metagenomic data with different methods.

## Discussion

Based on the information from the PUB water supply (https://www.pub.gov.sg/watersupply) there are four main “national taps” of water in Singapore, including water from the local catchment, water imported, highly-purified reclaimed water known as NEWater, the desalinated seawater. The local water catchment refers to the underground rainwater collection system and the reservoirs. The water imported will also be stocked at the reservoir before treated for human consumption. From these two points, reservoir water plays an essential role in the freshwater supply in Singapore. This is also the initial motivation to build a completely separate system for pollution, to protect the reservoir water quality better. Besides, the Singapore government generously encourages people to get close to the natural environment, many reservoirs like Chinese Garden, Pougle were built with the population entertainment functions. This increases the anthropogenic influence on the aquatic environment and, on the other hand, also increases the exposure of humans to microorganisms from the aquatic environment. As reported by Laurence Haller, et al previously, the wastewater from hospitals contains many types of CRE and ESBL carrier and genes(Haller, Chen et al. 2018), even though the rainwater collection is a separate system, there is still raise the concert if the reservoir waters have been contaminated with dangerous AMR pathogens. Unfortunately, the same ST isolates collected from different locations regardless of sources and time, in some way supported the hypothesis.

Refer to the current situation, in which the same ST resistant bacteria can be detected in all three sources in “One Health” approach, a genetic-based risk assessment system from the environment to humans is in urgent need. The present research is an example to process the first step of risk assessment as “hazard identification”, which refer to understand the potential risk bacteria to human health and also their concentrations. Even though the estimation is not working as expected, it still provided a brief idea for the upline of concentration. For the genetic-based risk assessment model, the non-pathogenetic bacteria which can transfer the AMR genes as a “vector” is also included. From our analysis, different species may play a different role in this model. The difference between *E. coli* and *Klebsiella pneumoniae* can give us some new sight of their different fate in AMR spreading. Based on the phylogenetic analysis with the same species from different sources, and the transferable beta-lactamase gene analysis, the *E. coli* from the reservoir has shown some genome-or isolate-level relatedness to the isolates from humans and food, as the same sequence type found among different sources. However, refer to *Klebsiella pneumoniae*, the isolates from different groups rarely shared the same ST. Instead, the same combination of resistance genes has been found among different isolates, and also, we have proved their capability to pass beta-lactamase to isolates to other species. From the limit data, the *E .coli* is performed more like a direct risk and spread at the genome/isolate-level, relatively, *Klebsiella spp*. perform more like a “vector” to contribute the resistance gene instead of colonized at different sources directly. However, all these hypotheses were based on the limited data we got, a national level of isolates collection and surveillance is needed to build the risk assessment system and generate a more accurate conclusion. The exposure model for contacting reservoir water is also missing, and more studies are expected. Also, at that stage, we can complete the source distribution with more data support to find AMR’s main sources for humans.

The prediction of AMR with genetic data has become a hot topic(Feldgarden, Brover et al. 2019). The new landed version of ResFinder has provided this function(Bortolaia, Kaas et al. 2020). However, the disagreement between phenotypic and genotypic addressed in this study has already been widely discussed in AMR detection(Hazards, Koutsoumanis et al. 2019). As mentioned previously, the novel mutation may change the enzymatic activity and influences the substrate type could be hydrolyzed, further affecting the resistance phenotype(Palzkill 2018). Therefore, further study for protein structure and function would be needed in the future. With more variants documented, the detection accuracy would be increased. Based on all the isolates we collected and their MIC result, we found three factors, which may influence the resistance pattern. First is the variant type of beta-lactamase. The mutation on the known resistance genes may change their resistance pattern, such as the mutation of *bla*_CTX-M-151_ found in our isolates, which may block their sensitivity to beta-lactamase inhibitor, and reduce their resistance to cephalosporin. The second is the genetic environment of the resistance gene. As the different distance of *bla*_CTX-M_ to the insert sequence has been found in our isolates, the distance to the upside gene and promotor is also varied, which may influence the transcription level regulation. The third is the species of the isolates. Significantly, both the *Klebsiella* spp. and *E. coli* isolates contain the same gene cluster of *bla*_CTX-M-15_, while their susceptibility is different. This phenome has also been observed as the different susceptibility between donors and transconjugants, who contain the same gene cluster. The regulation of beta-lactamase and how it would influence phenotypic resistance would be an interesting topic for further studies.

The conversed gene cluster reported here has been reported in ESBL-producing *E. coli* worldwide(Karim, Poirel et al. 2001, Dhanji, Patel et al. 2011). Interestingly, this time, we further proved it could also be detected on the plasmid of *Klebsiella* spp. The conjugation test has approved the transferability of the *bla*_CTX-M_ gene, which may suggest the *bla*_CTX-M_ genes are spreading in this conserved formate in Singapore. This gene cluster could become a potential indicator to evaluate the spreading of ESBL genes between different species.

The resistance genes detected from isolates and metagenomic annotation are quite different. There has been an ongoing discussion for the comparison between these two methods. However, as our research’s primary focus is the beta-lactamase genes, the shotgun sequencing may not be specific enough for this research. Other studies reported alluding to the need to enrich before sequencing, increasing the possibility of beta-lactamase detection(Noyes, Weinroth et al. 2017). But during the enrichment progress, as the growth rate for different species is different, some types of bacteria will become dominant, which will change the relative abundance of the resistance genes. From this angle, important quantification information is missing. An ideal case is to find a beta-lactamase gene as an indicator so that an amplicon metagenomic sequencing could be used to specifically target it. And this gene could be used to represent or evaluate the complete AMR spreading and would be useful for the AMR risk assessment.

There are still some limitations of this study, which can be further improved in a better setting in the future. Firstly, the missing estimation targeting specific species. In this study, our estimate focuses on the ESBL-carrier regardless of species, which lacks the prevalence data of ESBL-producing strain among certain species. For example, many studies reported the positive rate of ESBL-carrier among the total *E .coli* population, which didn’t present in this study. Besides, the estimation of the ESBL-carrier is not working as expected; MPN based methods may be applied to improve the accuracy for estimation. Secondly, the sample size is not big enough. To have a general idea of the AMR spreading, we collected the water sample as one reservoir per point instead of a multi-point collection for each reservoir. This is not just due to the limited manpower but also borrowed the idea of the outbreak investigation. Thirdly, we didn’t pick a significant mathematic model for tracking the AMR spreading pathway among different sources. Instead, this paper focuses on investigating the transferable gene cluster and the similarity of the genetic environment of beta-lactamase genes from various sources.

In conclusion, we have detected ESBL and carbapenem-resistant isolates, including the ESBL pathogens in reservoir water, which may suggest an anthropogenic influence on reservoirs. We also present the distributed spreading of AMR genes among different sources and different species. Besides, we found the conserved gene cluster and proved its transferability among different species, which may become an AMR spreading indicator in the future.

## Supporting information

Supplymentary_all

## Acknowledgment

The author would like to thank Dr. Jianming Zeng (University of Macau), Mr. Jiadong Zhao (Shenyou Genome Research Center at Nanjing) for their thoughtful suggestions on metagenomic sequencing analysis.

## Funding

The research was supported by Nanyang Technological University.

## Declaration

All authors declare no conflict of interests.

## Notes

### Competing Interest Statement

The authors have declared no competing interest.

