## Supplementary material for "Reservoir water in Singapore contains ESBL-producing and carbapenem-resistant bacteria with conjugatable conserved gene cluster transfer between different species": Supplymentary_all

**Table S1. Basic information of the isolates.**

| **Isolates ID** | **Reservoir ID** | **Species** | **MLST type** |
| --- | --- | --- | --- |
| BB10013-Y | Bedok | *Pseudomonas* spp. | N.A. |
| BedokY-2 | Bedok | *Pseudomonas* spp. | N.A. |
| BrichB-2Y | Bedok | *Pseudomonas putida* | 17 |
| BrichC-1Y | Bedok | *Pseudomonas putida* | 34* |
| BrichC-3Y | Bedok | *Pseudomonas putida* | 67* |
| BrichZ-10 | Bedok | *Stenotrophomonas maltophilia* | 390* |
| BrichZ-7 | Bedok | *Stenotrophomonas maltophilia* | 332* |
| LsrichE-8G | Lower Seletar | *Aeromonas hydrophila* | 230*,180* |
| LsrichA-4Y | Lower Seletar | *Citrobacter freundii* | N.A. |
| Lsblue-1 | Lower Seletar | *E. coli* | 1011 |
| Lsblue-2 | Lower Seletar | *E. coli* | 38 |
| LSB1003-2N | Lower Seletar | *Pseudomonas putida* | 36* |
| Lsrich E-2B | Lower Seletar | *Pseudomonas mosselii* | N.A. |
| LSB1002-1N | Lower Seletar | *Pseudomonas* spp*.* | N.A. |
| MrichA-1Y | Marina Bay | *Aeromonas enteropelogenes* | 310*,472*,285* |
| MarinaQ1 | Marina Bay | *E. coli* | 1193 |
| MarinaZ3 | Marina Bay | *Acinetobacter* spp. | N.A. |
| MarinaZ4 | Marina Bay | *Acinetobacter baumannii* | 1568 |
| MrichA-2Y | Marina Bay | *E. coli* | 206 |
| MaA1002-3Y | Marina Bay | *Bacillus pumilus* | N.A. |
| MaB1001-1B | Marina Bay | *Aeromonas caviae* | 310*,472*,285* |
| PrichA-15 | Punggol | *Aeromonas hydrophila* | 313 |
| **Isolates ID** | **Reservoir ID** | **Species** | **MLST type** |
| PrichA-3Y | Punggol | *E. coli* | 5276 |
| PA1012-1N | Punggol | *Pseudomonas alcaligenes* | N.A. |
| PA1013-1Y | Punggol | *Pseudomonas* spp. | N.A. |
| Prich D-1B | Punggol | *Pseudomonas mosselii* | N.A. |
| Prich7-2 | Punggol | *Aeromonas hydrophila* | 230*,232* |
| PrichB-4 | Punggol | *Klebsiella pneumoniae* | 441 |
| Srich E-5G1 | Serangoon | *Pseudomonas putida* | 82*,93* |
| Srich E-5G2 | Serangoon | *Pseudomonas mosselii* | N.A. |
| SrichD-2-2 | Serangoon | *Pseudomonas mosselii* | N.A. |
| SrichA-1 | Serangoon | *E. coli* | 410 |
| SA10126N | Serangoon | *Pseudomonas* spp. | N.A. |
| SblueE-7 | Serangoon | *E. coli* | 1642 |
| SblueE-3 | Serangoon | *E. coli* | 155 |
| SrichE-2G | Serangoon | *Aeromonas* spp. | 185* |
| SrichB-1 | Serangoon | *Klebsiella pneumoniae* | 1427 |
| CGA-4 | Chinese Garden | *Burkholderia cepacia* | 365*,1849*,1611* |
| CGB6 | Chinese Garden | *Chromobacterium violaceum* | N.A. |
| CGB1 | Chinese Garden | *Chromobacterium violaceum* | N.A. |
| CGA-5 | Chinese Garden | *Ochrobactrum* sp. | N.A. |
| CGX1 | Chinese Garden | *E. coli* | 1193 |
| CGX2 | Chinese Garden | *Klebsiella pneumoniae* | 194 |
| CGA-6 | Chinese Garden | *Ochrobactrum* sp. | N.A. |

*: alleles with less than 100% identity found, the nearest ST indicated. N.A.:Not included in the database.

**Table S2. MIC result of isolates.**

|  | Ceftriaxone | Meropenem | Cephalothin | Cefpodoxime | Ciprofloxacin | Cefotaxime | Cefotaxime / clavulanic acid |
| --- | --- | --- | --- | --- | --- | --- | --- |
| BB10013-Y | 8 | <1 | >16 | >32 | <1 | 8 | 8/4 |
| BedokY-2 | 16 | >8 | >16 | >32 | <1 | 16 | 8/4 |
| BrichZ-10 | 32 | >8 | >16 | >32 | <1 | 32 | 2/4 |
| BrichZ-7 | 128 | >8 | >16 | >32 | 2 | 64 | 16/4 |
| LsrichE-8G | 64 | <1 | >16 | >32 | <1 | 64 | 32/4 |
| LsrichA-4Y | 8 | <1 | >16 | >32 | <1 | 8 | 4/4 |
| Lsblue-1 | >128 | <1 | >16 | >32 | >2 | >64 | 0.12/4 |
| Lsblue-2 | >128 | <1 | >16 | >32 | <1 | >64 | 0.12/4 |
| LSB1002-1N | 16 | <1 | >16 | >32 | <1 | 16 | 16/4 |
| MrichA-1Y | >128 | <1 | >16 | >32 | >2 | >64 | 0.12/4 |
| MarinaQ1 | >128 | <1 | >16 | >32 | >2 | >64 | 0.12/4 |
| MarinaZ3 | 16 | <1 | >16 | 16 | <1 | 16 | 8/4 |
| MarinaZ4 | 16 | <1 | >16 | 16 | <1 | 16 | 16/4 |
| MrichA-2Y | >128 | <1 | >16 | >32 | >2 | >64 | 1/4 |
| MaB1001-1B | >128 | <1 | >16 | >32 | >2 | 64 | 0.12/4 |
| PrichA-15 | >128 | <1 | >16 | >32 | >2 | >64 | 1/4 |
| PrichA-3Y | 128 | <1 | >16 | >32 | <1 | 64 | 0.12/4 |
| PA1012-1N | >128 | >8 | >16 | >32 | <1 | >64 | >64/4 |
| Prich7-2 | 128 | <1 | >16 | >32 | >2 | 64 | <0.12/4 |
| PrichB-4 | 64 | <1 | >16 | >32 | 2 | 64 | <0.12/4 |
| SA10126N | 64 | 4 | >16 | >32 | <1 | 32 | 32/4 |
| SblueE-7 | >128 | <1 | >16 | >32 | <1 | >64 | <0.12/4 |
| SblueE-3 | >128 | <1 | >16 | >32 | <1 | >64 | <0.12/4 |
| SrichE-2G | 128 | <1 | >16 | >32 | <1 | 64 | 0.5/4 |
| SrichB-1 | >128 | <1 | >16 | >32 | >2 | >64 | <0.12/4 |
| CGX1 | >128 | <1 | >16 | >32 | >2 | >64 | <0.12/4 |
| CGX2 | 128 | <1 | >16 | >32 | 2 | 64 | <0.12/4 |
| CGA-6 | >128 | <1 | >16 | >32 | <1 | >64 | >64/4 |
| CGA-5 | >128 | >8 | >16 | >32 | <1 | >64 | >64\4 |

|  | Gentamicin | Ampicillin | Ceftazidime | Ceftazidime / clavulanic acid | Cefazolin | Imipenem | Piperacillin / tazobactam constant 4 |
| --- | --- | --- | --- | --- | --- | --- | --- |
| BB10013-Y | <4 | >16 | 1 | 1/4 | >16 | 1 | <4/4 |
| BedokY-2 | <4 | >16 | 2 | 2/4 | >16 | 4 | 16/4 |
| BrichZ-10 | <4 | >16 | 2 | 2/4 | >16 | >16 | 8/4 |
| BrichZ-7 | 8 | >16 | 4 | 4/4 | >16 | >16 | 64/4 |
| LsrichE-8G | <4 | >16 | 32 | 16/4 | >16 | <0.5 | <4/4 |
| LsrichA-4Y | <4 | >16 | 4 | 2/4 | >16 | <0.5 | 8/4 |
| Lsblue-1 | >16 | >16 | 4 | 0.25/4 | >16 | <0.5 | <4/4 |
| Lsblue-2 | <4 | >16 | 8 | 0.25/4 | >16 | <0.5 | <4/4 |
| LSB1002-1N | <4 | >16 | 4 | 4/4 | >16 | <0.5 | 8/4 |
| MrichA-1Y | <4 | >16 | >128 | 1/4 | >16 | <0.5 | 8/4 |
| MarinaQ1 | >16 | >16 | 8 | 0.25/4 | <8 | <0.5 | <4/4 |
| MarinaZ3 | <4 | >16 | 8 | 8/4 | >16 | 1 | 8/4 |
| MarinaZ4 | <4 | >16 | 8 | 8/4 | >16 | <0.5 | 32/4 |
| MrichA-2Y | 16 | >16 | >128 | 1/4 | >16 | <0.5 | 16/4 |
| MaB1001-1B | <4 | >16 | >128 | 0.12/4 | >16 | <0.5 | <4/4 |
| PrichA-15 | 16 | >16 | >128 | 0.5/4 | >16 | <0.5 | >64/4 |
| PrichA-3Y | <4 | >16 | 4 | 0.12/4 | >16 | <0.5 | <4/4 |
| PA1012-1N | <4 | >16 | 64 | 64/4 | >16 | 16 | >64/4 |
| Prich7-2 | >16 | >16 | 16 | 0.25/4 | >16 | <0.5 | 32/4 |
| PrichB-4 | >16 | >16 | 32 | 0.5/4 | >16 | <0.5 | 16/4 |
| SA10126N | <4 | >16 | 16 | 16/4 | >16 | 2 | 32/4 |
| SblueE-7 | <4 | >16 | 16 | 0.25/4 | >16 | <0.5 | <4/4 |
| SblueE-3 | <4 | >16 | 16 | <0.12/4 | >16 | <0.5 | <4/4 |
| SrichE-2G | <4 | >16 | 4 | 1/4 | >16 | <0.5 | <4/4 |
| SrichB-1 | >16 | >16 | 32 | 0.25/4 | >16 | 1 | 16/4 |
| CGX1 | >16 | >16 | 4 | <0.12/4 | >16 | <0.5 | <4/4 |
| CGX2 | <4 | >16 | 32 | 0.5/4 | >16 | <0.5 | 16/4 |
| CGA-6 | 8 | >16 | >128 | >128/4 | >16 | 4 | >64/4 |
| CGA-5 | <4 | >16 | >128 | >128\4 | >16 | 1 | >64\4 |

|  | Cefepime | Cefoxitin |
| --- | --- | --- |
| BB10013-Y | <1 | >64 |
| BedokY-2 | <1 | >64 |
| BrichZ-10 | 8 | >64 |
| BrichZ-7 | 16 | >64 |
| LsrichE-8G | 8 | 8 |
| LsrichA-4Y | <1 | >64 |
| Lsblue-1 | 16 | 8 |
| Lsblue-2 | >16 | 8 |
| LSB1002-1N | 4 | >64 |
| MrichA-1Y | >16 | 16 |
| MarinaQ1 | >16 | <4 |
| MarinaZ3 | 4 | >64 |
| MarinaZ4 | 4 | >64 |
| MrichA-2Y | >16 | 32 |
| MaB1001-1B | 8 | <4 |
| PrichA-15 | >16 | 16 |
| PrichA-3Y | 4 | <4 |
| PA1012-1N | >16 | >64 |
| Prich7-2 | 8 | 8 |
| PrichB-4 | 8 | <4 |
| SA10126N | 4 | >64 |
| SblueE-7 | >16 | 8 |
| SblueE-3 | >16 | <4 |
| SrichE-2G | 8 | <4 |
| SrichB-1 | 8 | <4 |
| CGX1 | 16 | <4 |
| CGX2 | 8 | <4 |
| CGA-6 | >16 | >64 |
| CGA-5 | >16 | >64 |

>: over the detection range. <: start to grow at the lowest concentration

| **Isolate information** | | **Replicon information** | **Contigs information** | | |
| --- | --- | --- | --- | --- | --- |
| **Isolate ID** | **Species** | **Ori type** | **Contigs ID** | **Contigs length/bp** | **Coverage** |
| Lsblue-1 | *E. coli* | Col(BS512) | 55 | 2101 | 594.6 |
|  |  | IncFIB(AP001918)* | 41 | 10127 | 84.8 |
|  |  | IncFII(pCoo)* | 4 | 60476 | 93.1 |
|  |  | IncI(Gamma)* | 13 | 20644 | 101.5 |
| Lsblue-2 | *E. coli* | IncFII(pHN7A8)* | 3 | 70476 | 104.8 |
| MarinaQ1 | *E. coli* | Col(BS512) | 61 | 2113 | 1682.3 |
|  |  | Col156* | 19 | 10578 | 151 |
|  |  | IncFIA* | 6 | 27711 | 163.8 |
|  |  | IncFIB(AP001918)* | 26 | 12377 | 142.2 |
|  |  | IncI1-I(Gamma) | 24 | 20948 | 187.4 |
| MrichA-2Y | *E. coli* | ColE10 | 6 | 10566 | 1515.9 |
|  |  | IncI1-I(Gamma) | 31 | 28462 | 156 |
| PrichA-3Y | *E. coli* | IncFII(pCoo)* | 61 | 9158 | 133.7 |
|  |  | IncI1-I(Gamma)* | 68 | 17242 | 118.4 |
|  |  | p0111* | 39 | 12487 | 74.8 |
| SrichA-1 | *E. coli* | Col(BS512) | 24 | 2088 | 293.1 |
|  |  | IncFIA* | 36 | 31098 | 98.1 |
|  |  | IncFIB(AP001918)* | 53 | 13204 | 92.3 |
|  |  | IncFII(pAMA1167-NDM-5) | 27 | 5981 | 98.3 |
|  |  | IncI2(Delta)* | 22 | 38151 | 104.8 |
| SblueE-7 | *E. coli* | IncFIA(Hl1)* | 40 | 7802 | 184.26 |
|  |  | IncFIB(K)* | 34 | 11336 | 111.95 |
|  |  | IncR* | 15 | 9096 | 154.37 |
|  |  | IncY* | 26 | 7497 | 116.8 |
| CGX1 | *E. coli* | Col(BS512) | 1 | 2113 | 706.33 |
|  |  | Col(MG828)* | 4 | 1552 | 1119.92 |
|  |  | Col156* | 56 | 10578 | 87 |
|  |  | IncFIA* | 17 | 27711 | 105.49 |
|  |  | IncFIB(AP001918)* | 37 | 12377 | 81.8 |
|  |  | IncI1-I(Gamma)* | 18 | 20948 | 98.97 |
| CGX2 | *Klebsiella pneumoniae* | IncFIB(K)* | 16 | 34145 | 79.5 |
|  |  | IncFII(K)* | 2 | 43684 | 82.1 |
| PrichB-4 | *Klebsiella pneumoniae* | IncFIB(K)* | 34 | 9934 | 80.6 |
|  |  | IncFIB(pKPHS1)* | 12 | 109651 | 100 |
|  |  | IncFII(K)* | 3 | 58405 | 83.4 |
| SrichB-1 | *Klebsiella pneumoniae* | IncFIB(K)* | 29 | 74843 | 82.1 |

**Table S3.** Ori detected with PlasmidFinder. * indicate the one without 100% identification.

**Table S4.** **Best hits of contigs carrying AMR genes found by Blastn.**

| **Isolate ID** | **Resistance information** | | **Contigs information** | | | **Best hits found by Blastn** | | | | | | | | |
| --- | --- | --- | --- | --- | --- | --- | --- | --- | --- | --- | --- | --- | --- | --- |
|  | **Beta-lactamase Class** | **Beta-lactamase gene** | **Contigs ID** | **Contigs length/bp** | **Coverange** | **Reference Genome** | **Reference plasmid** | **Location** | **Accession** | **Source/Country** | **Total score** | **Query cover** | **E value** | **Identity** |
| BB10013-Y | B | *bla*_PAM-1*_ | 13 | 280071 | 53.7 | *Pseudomonas citronellolis* strain SJTE-3 | N.A. | Chromosome | NZ_CP015878 | Sludge/China | 7.80E+05 | 70% | 0 | 86.37% |
| BedokY-2 | B | *bla*_POM-1*_ | 10 | 408960 | 63 | *Pseudomonas citronellolis* strain SJTE-3 | N.A. | Chromosome | NZ_CP015878 | Sludge/China | 5.11E+05 | 54% | 0 | 81.13% |
| PA1012-1N | B | *bla*_PAM-1*_ | 8 | 246341 | 126.5 | *Pseudomonas citronellolis* strain SJTE-3 | N.A. | Chromosome | NZ_CP015878 | Sludge/China | 3.92E+05 | 48% | 0 | 79.64% |
| SA10126N | B | *bla*_PAM-1*_ | 19 | 54236 | 61.3 | *Pseudomonas sp.* DY-1 | N.A. | Chromosome | CP032616 | Soil/China | 18637 | 30% | 0 | 86.30% |
| LSB1002-1N | D | *bla*_OXA-457*_ | 6 | 451899 | 52.1 | *Pseudomonas nitroreducens* strain HBP1 | N.A. | Chromosome | NZ_CP049140 | Wastewater/ Switzerland | 1.20E+06 | 95% | 0 | 99.94% |
| BrichZ-10 | B | *bla*_L1*_ | 42 | 821217 | 82.6 | *Stenotrophomonas maltophilia* strain FDAARGOS_325 | N.A. | Chromosome | NZ_CP022053 | Human/USA | 3.02E+06 | 86% | 0 | 92.82% |
| BrichZ-7 | B | *bla*_L1*_ | 12 | 783456 | 94 | *Stenotrophomonas maltophilia* strain CSM2 | N.A. | Chromosome | NZ_CP025298 | Laboratory sink/ Mexico | 2.91E+06 | 94% | 0 | 92.36% |
| LsrichE-8G | C | *bla*_MOX-5*_ | 123 | 22335 | 66.4 | *Aeromonas sp.* ASNIH5 | N.A. | Chromosome | CP026122 | Wastewater/USA | 48481 | 99% | 0 | 98.37% |
|  | D | *bla*_OXA-427*_ | 2 | 81891 | 63.3 | *Aeromonas caviae* strain 8LM | N.A. | Chromosome | CP024198 | Human/Brazil | 1.35E+05 | 90% | 0 | 98.70% |
| MrichA-1Y | A | *bla*_PER-3_ | 31 | 7992 | 44.3 | *Aeromonas hydrophila* strain RJ604 | N.A | Integron | KU133344 | Human/China | 15422 | 100% | 0 | 99.99% |
|  | C | *bla*_MOX-5*_ | 324 | 206386 | 50.7 | *Aeromonas caviae* strain WCW1-2 | N.A. | Chromosome | CP039832 | Sewage/China | 3.35E+05 | 80% | 0 | 99.38% |
|  | D | *bla*_OXA-427*_ | 34 | 123786 | 52.8 | *Aeromonas caviae* strain WCW1-2 | N.A. | Chromosome | CP039832 | Sewage/China | 1.93E+05 | 77% | 0 | 99.27% |
| MaB1001-1B | D | *bla*_OXA-427*_ | 37 | 100720 | 61.5 | *Aeromonas caviae* strain WCW1-2 | N.A. | Chromosome | CP039832 | Sewage/China | 1.79E+05 | 88% | 0 | 99.66% |
|  | C | *bla*_MOX-5*_ | 122 | 206936 | 66.8 | *Aeromonas caviae* strain WCW1-2 | N.A. | Chromosome | CP039832 | Sewage/China | 3.40E+05 | 80% | 0 | 99.38% |
| PrichA-15 | D | *bla*_OXA-10_ | 431 | 57028 | 59 | *Aeromonas caviae strain* WCW1-2 | N.A. | Chromosome | CP039832 | Sewage/China | 1.106e+05 | 80% | 0 | 98.93% |
|  | A | *bla*_VEB-1*_ |  |  |  |  |  |  |  |  |  |  |  |  |
|  | D | *bla*_OXA-427*_ | 88 | 35361 | 64.2 | *Aeromonas caviae strain* R25-2 | N.A. | Chromosome | CP025777 | Wastewater/China | 62113 | 100% | 0 | 98.02% |
|  | C | *bla*_MOX-2_ | 56 | 52941 | 64.3 | *Aeromonas sp.* ASNIH1 | N.A. | Chromosome | CP026228 | Wastewater/USA | 97762 | 99% | 0 | 98.31% |
| Prich7-2 | D | *bla*_OXA-427*_ | 21 | 35379 | 52.4 | *Aeromonas caviae strain* NCTC12244 | N.A. | Chromosome | LS483441 | N.A./UK | 61934 | 96% | 0 | 97.49% |
|  | A | *bla*_PER-3_ | 149 | 4798 | 67.8 | *Aeromonas hydrophila* strain RJ604 | N.A | Integron | KU133344 | Human/China | 8987 | 97% | 0 | 100% |
|  | C | *bla*_MOX-6*_ | 799 | 28454 | 47.7 | *Aeromonas caviae* strain T25-39 | N.A. | Chromosome | CP025706 | Wastewater/China | 61164 | 99% | 0 | 99.10% |
|  | A | *bla*_TEM-1B_ | 49 | 47258 | 80.2 | *Aeromonas sp.* ASNIH4 | pAER-f909 | Plasmid | CP026221 | Wastewater/USA | 58158 | 84% | 0 | 90.93% |
| SrichE-2G | D | *bla*_OXA-427*_ | 9 | 114376 | 71.3 | *Aeromonas sp.* ASNIH1 | N.A. | Chromosome | CP026228 | Wastewater/USA | 1.46E+05 | 87% | 0 | 91.93% |
|  | A | *bla*_CTX-M-3_ | 5 | 226087 | 70.4 | *Aeromonas sp.* ASNIH1 | N.A. | Chromosome | CP026228 | Wastewater/USA | 3.34E+05 | 80% | 0 | 91.19% |
|  | C | *bla*_MOX-6*_ | 4 | 1508227 | 69.4 | *Aeromonas sp.* ASNIH1 | N.A. | Chromosome | CP026228 | Wastewater/USA | 1.94E+06 | 86% | 0 | 90.40% |

| **Isolate ID** | **Resistance information** | | **Contigs information** | | | **Best hits found by NCBI Blastn** | | | | | | | | |
| --- | --- | --- | --- | --- | --- | --- | --- | --- | --- | --- | --- | --- | --- | --- |
|  | **Beta-lactamase Class** | **Beta-lactamase gene** | **Contigs ID** | **Contigs length/bp** | **Coverange** | **Reference Genome** | **Reference plasmid** | **Location** | **Accession** | **Source/ Country** | **Total score** | **Query cover** | **E value** | **Identity** |
| LsrichA-4Y | A | *bla*_CTX-M-151*_ | 1 | 286113 | 117.7 | *Citrobacter freundii* CFNIH1 | N.A. | Chromosome | NZ_CP007557 | Sink aerator/ USA | 1.63E+05 | 73% | 0 | 82.74% |
| Lsblue-1 | A | *bla*_CTX-M-65_ | 32 | 4893 | 99.5 | *Salmonella enterica subsp. enterica serovar* SPE100 | pSPE100_vir | Plasmid | CP040064 | Human/ Mexico | 13314 | 100% | 0 | 100% |
|  | A | *bla*_TEM-1B*_ | 81 | 474 | 182.3 | *Klebsiella pneumoniae* strain BJ20 | pBJ20-KPC | Plasmid | MT108208 | N.A./China | 1278 | 100% | 0 | 100% |
| Lsblue-2 | A | *bla*_CTX-M-15_ | 38 | 289346 | 61.4 | *Escherichia coli* strain CFSAN061770 | N.A. | Chromosome | NZ_CP023142 | Food/Egypt | 4.39E+05 | 71% | 0 | 98.83% |
| MrichA-2Y | A | *bla*_CTX-M-55_ | 62 | 2795 | 189.2 | *Salmonella sp.* strain PJM1 | pPJM1 | Plasmid | MN539017 | Chicken/ China | 5432 | 100% | 0 | 100% |
|  | A | *bla*_TEM-1C_ | 31 | 28462 | 156 | *Escherichia coli* strain Ec-050 | pEc-050-TEM-30 | Plasmid | CP043228 | Human/ Switzerland | 50519 | 95% | 0 | 99.95% |
| MarinaQ1 | A | *bla*_TEM-1B_ | 51 | 1387 | 149 | *Escherichia coli* strain 5P | pVSI_NDM_5 | Plasmid | MN197360 | Human/ Czechia | 7541 | 100% | 0 | 100% |
|  | A | *bla*_CTX-M-15_ | 116 | 220574 | 63.8 | *Escherichia coli* strain MCJCHV-1 | N.A. | Chromosome | NZ_CP030111 | Human/USA | 4.91E+05 | 97% | 0 | 100% |
| PrichA-3Y | A | *bla*_CTX-M-15_ | 11 | 18365 | 116.5 | *Escherichia coli O169:H41* strain 2014EL-1345-2 | unnamed3 | Plasmid | CP024226 | N.A./USA | 350750 | 100% | 0 | 99.99% |
| SrichA-1 | C | *bla*_CMY-2_ | 52 | 22678 | 49.4 | *Escherichia coli* strain E118 | N.A. | Chromosome | CP049196 | Animal/ Korea | 85845 | 100% | 0 | 100% |
|  | B | *bla*_NDM-5_ | 29 | 5726 | 96.6 | *Escherichia coli* strain 52148 | p52148_NDM_5 | Plasmid | CP050384 | Human/ Czechia | 11291 | 100% | 0 | 99.98% |
|  | A | *bla*_CTX-M-15_ | 45 | 3967 | 98.2 | *Escherichia coli* isolate EcMAD1 | pEcMAD1 | Plasmid | LR595692 | Human/ Frace | 11901 | 100% | 0 | 100% |
|  | A | *bla*_TEM-1B_ | 101 | 1882 | 89.9 | *Escherichia coli* isolate EcMAD1 | pEcMAD1 | Plasmid | LR595692 | Human/ Frace | 10099 | 100% | 0 | 100% |
|  | D | *bla*_OXA-1_ | 85 | 2314 | 93.6 | *Vibrio cholerae* strain YA00120881 | pYA00120881 | Plasmid | MT151380 | Human/ Frace | 7468 | 100% | 0 | 100% |
| SblueE-7 | A | *bla*_CTX-M-55_ | 19 | 10162 | 87.1 | *Salmonella enterica subsp. enterica* strain CFSA1096 | pCFSA1096 | Plasmid | CP033347 | Food/China | 25620 | 100% | 0 | 100% |
| SblueE-3 | A | *bla*_CTX-M-55_ | 39 | 10213 | 102.5 | *Salmonella enterica subsp. enterica* strain CFSA1096 | pCFSA1096 | Plasmid | CP033347 | Food/China | 26709 | 100% | 0 | 100% |
| CGX1 | A | *bla*_TEM-1B_ | 31 | 1387 | 96.3 | *Escherichia coli* strain SCU-109 | pSCU-109-1 | Plasmid | CP051734 | Human/USA | 3166 | 100% | 0 | 100% |
|  | A | *bla*_CTX-M-15_ | 53 | 218263 | 43.9 | *Escherichia coli* strain MCJCHV-1 | N.A. | Chromosome | NZ_CP030111 | Human/USA | 4.86E+05 | 97% | 0 | 100% |
| PrichB-4 | A | *bla*_SHV-72*_ | 1 | 290180 | 52 | *Klebsiella pneumoniae* strain 203 | N.A. | Chromosome | NZ_CP021165 | Human/USA | 6.17E+05 | 99% | 0 | 99.98% |
|  | A | *bla*_TEM-1B_ | 32 | 8938 | 80.3 | *Klebsiella pneumoniae* strain E16KP0288 | pE16K0288-1 | Plasmid | CP052263 | Human/ Korea | 16854 | 100% | 0 | 100% |
|  | A | *bla*_CTX-M-15_ |  |  |  |  |  |  |  |  |  |  |  |  |
|  | D | *bla*_OXA-1_ | 48 | 2329 | 86 | *Klebsiella pneumoniae* strain E16KP0204 | pE16KP0204-1 | Plasmid | CP052298 | Human/ Korea | 8864 | 100% | 0 | 100% |

| **Isolate ID** | **Resistance information** | | **Contigs information** | | **Best hits found by NCBI Blastn** | | | | | | | | | |
| --- | --- | --- | --- | --- | --- | --- | --- | --- | --- | --- | --- | --- | --- | --- |
|  | **Beta-lactamase Class** | **Beta-lactamase gene** | **Contigs ID** | **Contigs length/bp** | **Coverange** | **Reference Genome** | **Reference plasmid** | **Location** | **Accession** | **Source/ Country** | **Total score** | **Query cover** | **E value** | **Identity** |
| SrichB-1 | A | *bla*_TEM-1B_ | 53 | 8882 | 71.4 | *Klebsiella pneumoniae* strain E16KP0288 | pE16K0288-1 | Plasmid | CP052263 | Human/Korea | 16647 | 100% | 0 | 100% |
|  | A | *bla*_CTX-M-15_ |  |  |  |  |  |  |  |  |  |  |  |  |
|  | D | *bla*_OXA-1_ | 112 | 2273 | 84.7 | *Klebsiella pneumoniae* strain E16KP0204 | pE16KP0204-1 | Plasmid | CP052298 | Human/Korea | 7003 | 100% | 0 | 100% |
|  | A | *bla*_SHV-40*_ | 130 | 511583 | 81.2 | *Klebsiella pneumoniae* strain INF249 | N.A. | Chromosome | NZ_CP024489 | Human/USA | 1.04E+06 | 97% | 0 | 99.19% |
| CGX2 | A | *bla*_TEM-1B_ | 14 | 11754 | 86.3 | *Klebsiella pneumoniae* strain E16KP0288 | pE16K0288-1 | Plasmid | CP052263 | Human/Korea | 23802 | 100% | 0 | 100% |
|  | A | *bla*_CTX-M-15_ |  |  |  |  |  |  |  |  |  |  |  |  |
|  | D | *bla*_OXA-1_ | 40 | 2330 | 80.1 | *Klebsiella pneumoniae* strain E16KP0204 | pE16KP0204-1 | Plasmid | CP052298 | Human/Korea | 8881 | 100% | 0 | 100% |
|  | A | *bla*_SHV-61_ | 35 | 1275886 | 56.1 | *Klebsiella pneumoniae* strain INF249 | N.A. | Chromosome | NZ_CP024489 | Human/USA | 2.47E+06 | 94% | 0 | 99.32% |
| MarinaZ3 | C | *bla*_ADC-25*_ | 32 | 589745 | 159.3 | *Acinetobacter pittii* PHEA-2 | N.A. | Chromosome | NC_016603 | Wastewater/USA | 9.94E+05 | 96% | 0 | 97.01% |
|  | D | *bla*_OXA-506*_ | 35 | 1111341 | 164.2 | *Acinetobacter pittii PHEA-2* | N.A. | Chromosome | NC_016603 | Wastewater/USA | 1.65E+06 | 85% | 0 | 95.96% |
| MarinaZ4 | C | *bla*_ADC-25*_ | 16 | 599676 | 81.3 | *Acinetobacter baumannii* strain XH858 | N.A. | Chromosome | NZ_CP014528 | Human/China | 9.90E+05 | 91% | 0 | 98.74% |
|  | D | *bla*_OXA-343_ | 21 | 394432 | 77 | *Acinetobacter baumannii* strain XH858 | N.A. | Chromosome | NZ_CP014528 | Human/China | 5.94E+05 | 84% | 0 | 98.49% |
| CGA-6 | C | *bla*_OCH-2*_ | 14 | 277306 | 73.5 | *Ochrobactrum anthropi* strain OAB | N.A. | Chromosome | NZ_CP008820 | N.A./USA | 3.59E+05 | 77% | 0 | 86.78% |
| CGA-5 | C | *bla*_OCH-7*_ | 40 | 1011436 | 84.4 | *Ochrobactrum anthropi* ATCC 49188 | N.A. | Chromosome | NC_009667 | Type strain | 1.27E+06 | 89% | 0 | 87.25% |

The hits with the highest total score were present. N.A. : source information is not available. *: without 100% identification.

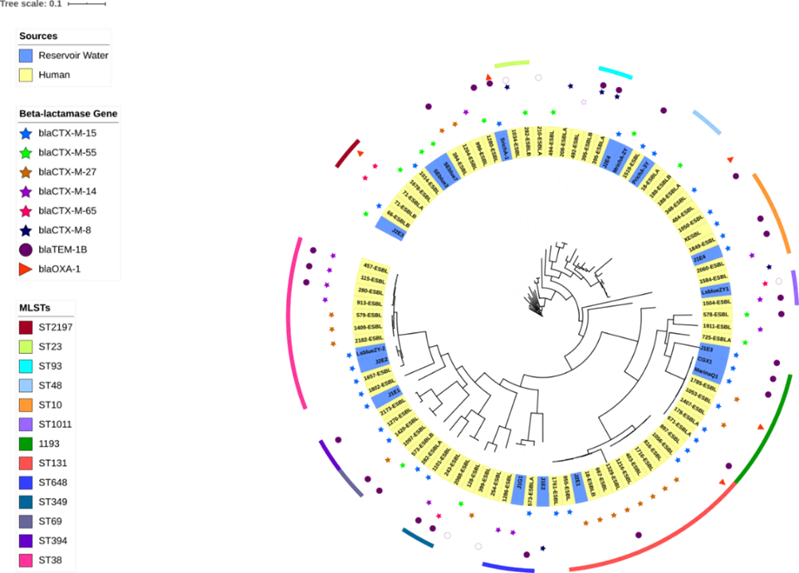

**Figure S1.** **Phylogenetic analysis of E.coli strain from reservoir water and healthy donors**. The phylogenetic tree was built based on core genome SNP, with MG1655 as reference. The inner-circle refers to the isolates' source, including 71 isolates from the health community and 18 from the reservoir water collected in this and previous research. The outer circle refers to the MLSTs. Only STs with more than two isolates were present.

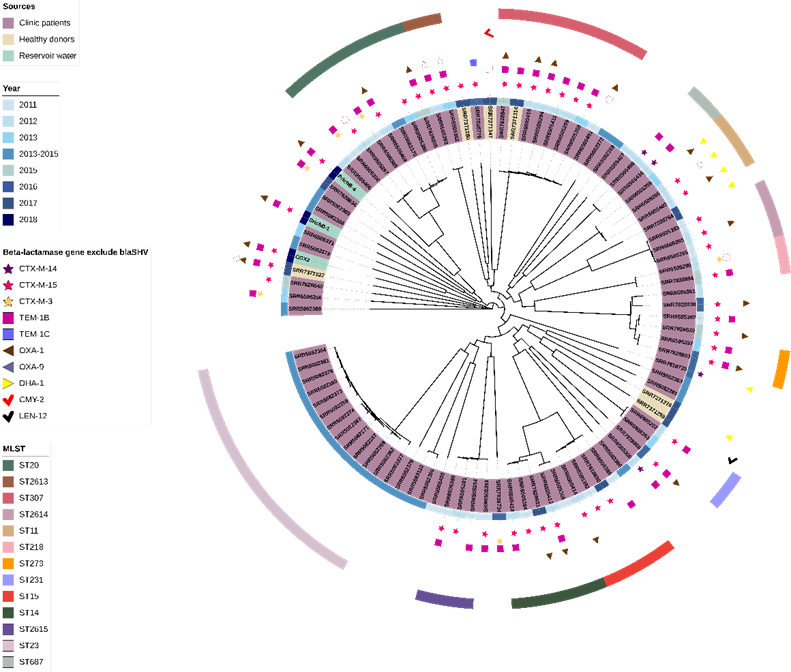

**Fig S2. Phylogenetic tree of *Klebsiella* spp. with beta-lactamase genes annotation.** Ninety-five isolates from four Bioproject, including strains from reservoir water, healthy donors, and clinic patients, have been plotted according to core genome SNP with strain HS11286 as the reference genome. The inner layer presents the isolation date (year), the outer layer presents the ST, only ST with more than three leaves was annotated. The beta-lactamases gene, except for *bla*SHV-like genes, were labeled in between the two-layer.

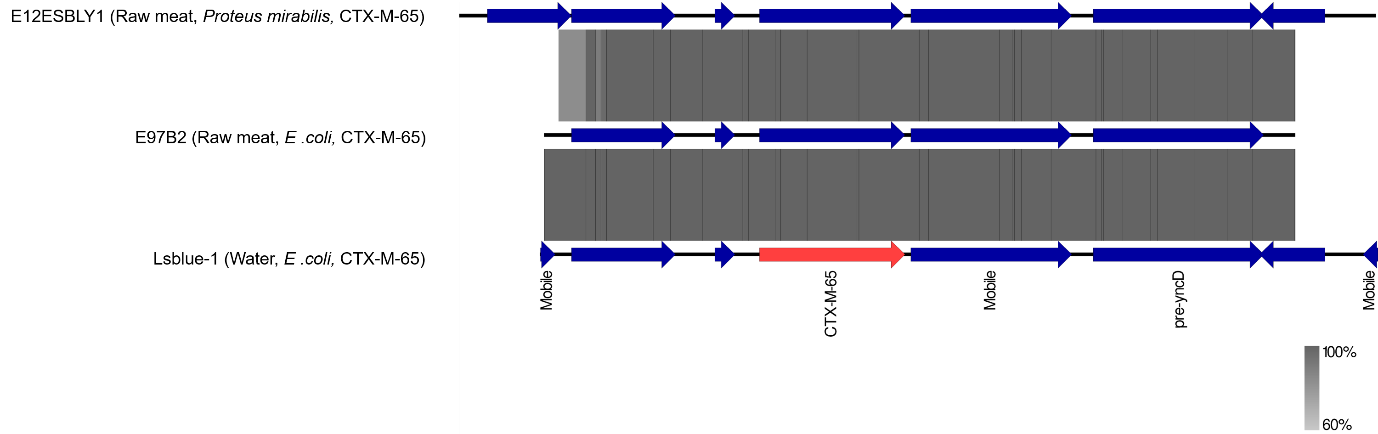

**Fig S3. Genetic environment analysis of CTX-M-65.** Contigs carrying CTX-65 of different species isolates, collected from multiple sources, have been aligned with Blast+. The high similarity among these contigs suggests the genetic environment of CTX-M-65 is highly conserved.

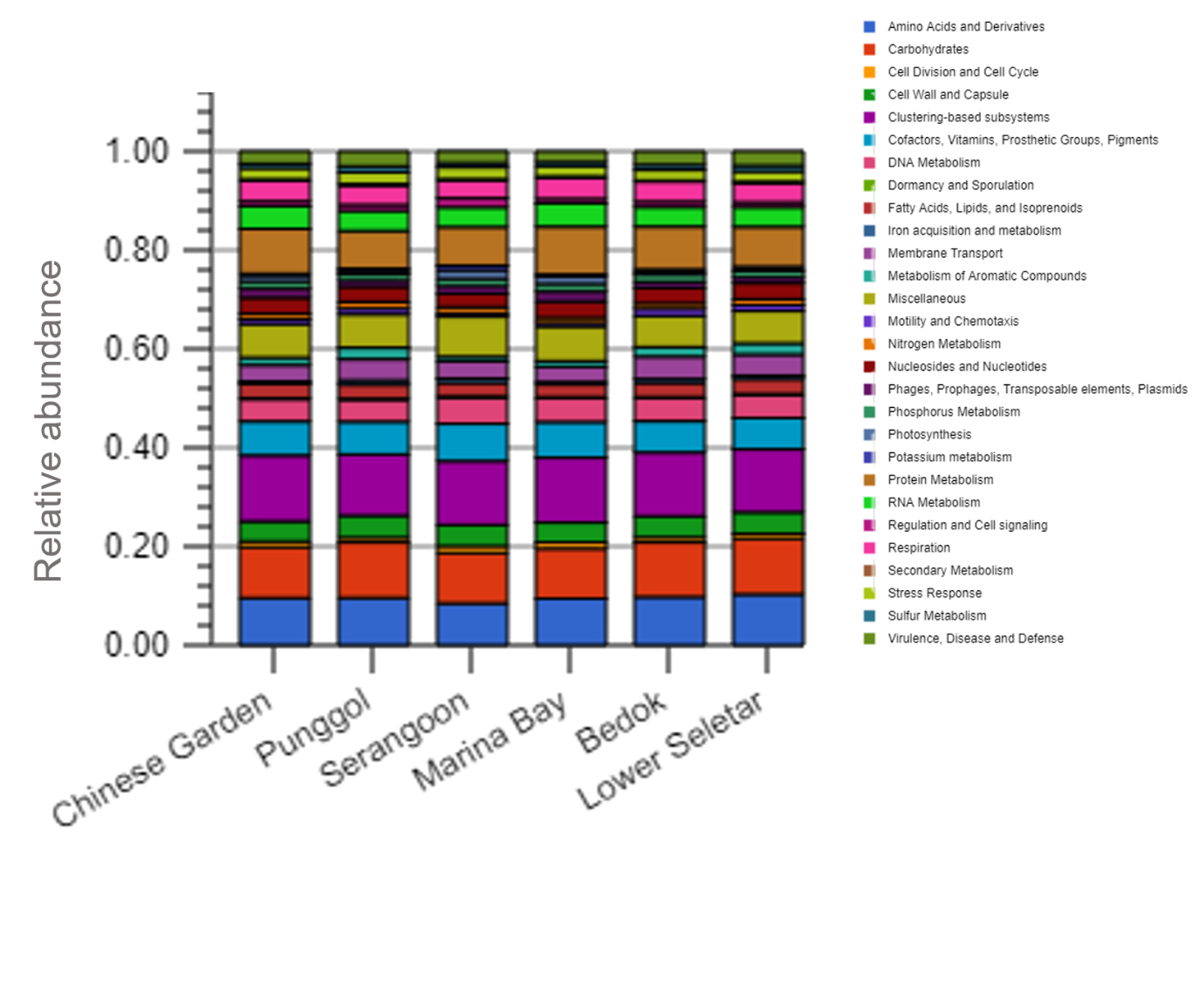

**Fig S4. Functional gene annotation of metagenomic data with MG-RAST.** The functional gene mapping was processed with MG-RAST and grouped according to the SEED database. The relative abundance was calculated based on the reading categorized into different functional groups and normalized to total functional readings.

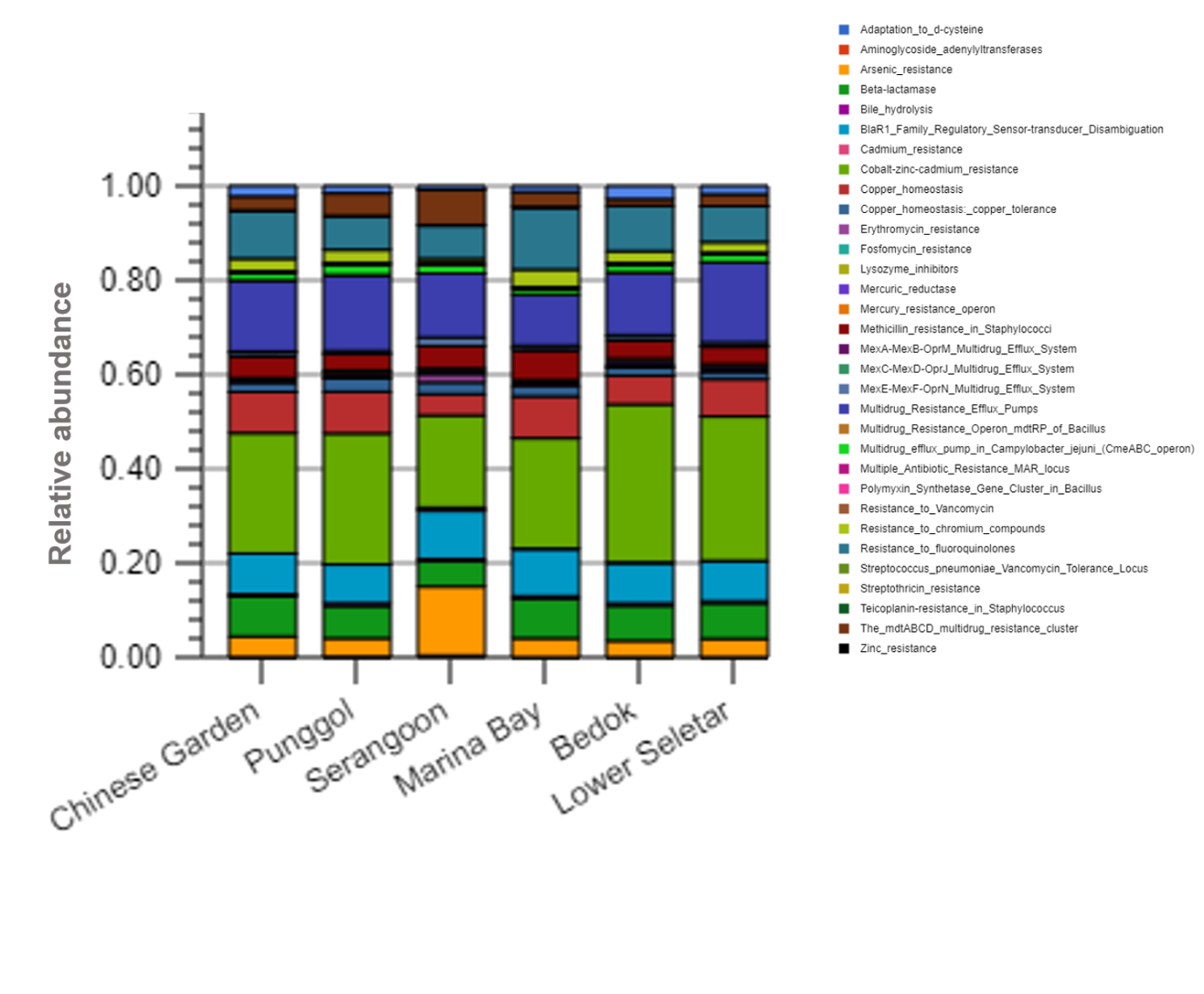

**Fig S5. Relative abundance of functional readings under the “Drug Resistance” catalog.** The relative abundance is calculated based on mapping reads normalized to total reads of the “Drug Resistance” catalog.
